# NOD2 and TLR2 Signal via TBK1 and PI31 to Direct Cross-presentation and CD8 T Cell Responses

**DOI:** 10.1101/570705

**Authors:** Daniele Corridoni, Seiji Shiraishi, Thomas Chapman, Tessa Steevels, Daniele Muraro, Marie-Laëtitia Thézénas, Gennaro Prota, Ji-Li Chen, Uzi Gileadi, Nicola Ternette, Vincenzo Cerundolo, Alison Simmons

**Author notes:** Corresponding Authors Alison Simmons; Daniele Corridoni.

## Abstract

NOD2 and TLR2 recognize components of bacterial cell wall peptidoglycan and direct defense against enteric pathogens. CD8^+^ T cells are important for immunity to such pathogens but how NOD2 and TLR2 induce antigen specific CD8^+^ T cell responses is unknown. Here, we define how these pattern recognition receptors (PRRs) signal in primary dendritic cells (DCs) to influence MHC class I antigen presentation. We show NOD2 and TLR2 phosphorylate PI31 via TBK1 following activation in DCs. PI31 interacts with TBK1 and Sec16A at endoplasmic reticulum exit sites (ERES), which positively regulates MHC class I peptide loading and immunoproteasome stability. Following NOD2 and TLR2 stimulation, depletion of PI31 or inhibition of TBK1 activity *in vivo* impairs DC cross-presentation and CD8^+^ T cell activation. DCs from Crohn’s patients expressing *NOD2* polymorphisms show dysregulated cross-presentation and CD8^+^ T cell responses. Our findings reveal unidentified mechanisms that underlie CD8^+^ T cell responses to bacteria in health and in Crohn’s.

## Introduction

NOD2 and TLR2 are pattern recognition receptors (PRRs) that recognize conserved components of peptidoglycan (PGN) found in bacterial cell walls (1,2). Both PRRs activate intracellular signaling pathways in antigen presenting cells (APCs) that drive pro-inflammatory and antimicrobial adaptive responses. Both receptors are important for clearance of bacterial pathogens such as *Salmonella typhimurium*, *Shigella flexneri* and *Mycobacterium tuberculosis* (3-5). Both NOD2 and TLR2 employ distinct signaling cascades to drive pro-inflammatory cytokine responses, however, the mechanism by which they connect to MHC class I antigen presentation machinery is unclear.

NOD2 cross-talks with TLR2 in myeloid cells. This manifests as amplification in signaling culminating in greatly heightened pro-inflammatory cytokine responses (6,7). Furthermore, gene expression studies have revealed dual activation of both receptors leads to a transcriptional programme that not only amplifies the differential expression common to both receptors, but leads to induction of a specific NOD2/TLR2 gene panel highlighting a physiological role for this cross-talk in differentiating pathogenic versus commensal invasion of APCs (8,7,9). NOD2 is the strongest associated Crohn’s susceptibility gene and Crohn’s disease patients who express NOD2 polymorphisms display loss of function for induction of NOD2 effector genes and NOD2/TLR2 specific genes (10,7,11).

NOD2 and TLR2 activate autophagy in myeloid cells to degrade invading bacteria and facilitate MHC class II antigen presentation (12-14). Stimulation of either receptor regulates Th1, Th2 and Th17 immune responses (15-18). DCs that have engulfed bacteria also present exogenous antigens on MHC I via cross-presentation, which is critical for CD8^+^ T cell responses against microbial pathogens (19). Proposed pathways and mechanisms underlying cross-presentation include the phagocytic pathway, which exports antigens from the phagosome to the cytosol, degradation by the immunoproteasome and peptide loading in the endoplasmic reticulum (ER) or in phagosomes (20). Alternatively, in cross-presentation through the vacuolar pathway, endosomal or phagosomal proteases degrade internalised antigens independent of immunoproteasomal degradation and loading MHC class I molecules occur in endocytic compartments (21).

PRR engagement increases CD8^+^ T cell activation by cross-presented peptides, but the molecular mechanisms underlying these effects are not completely defined (22,23,4,24,19,25,26). TLR4-dependent phosphorylation of phagosomal SNAP-23 recruits MHC class I molecules from endosomal recycling compartments to phagosomes, which promotes cross-presentation (27). However, the mechanisms by which either NOD2 or TLR2 signal to the MHC class I antigen presentation machinery to enhance CD8^+^ T cell activation remain unclear. We hypothesized undertaking an unbiased screen of NOD2 and TLR2 signaling in primary DCs would reveal molecules co-opted by these receptors to enable cross-presentation. Utilizing information obtained from a quantitative phosphoproteomic analysis comparing NOD2 and TLR2 signaling in human DCs, we discovered that NOD2 in combination with TLR2 phosphorylates PI31. This signaling pathway links PI31 with defective cross-presentation as found in Crohn’s.

## Materials and Methods

### Study Design

The objective of this study was to explore the role of NOD2 and TLR2 in cross-presentation in human dendritic cells undertaking an unbiased screen. We have used a quantitative phosphoproteomic analysis by liquid chromatography-tandem mass spectrometry (LC-MS/MS) followed by a computational analysis to identify the proteins as differentially abundant in response to NOD2 and TLR2 sensing. Validation of the phosphoproteomic analysis was performed by the detection of proteins in phosphoenriched lysates and detected by western blot. Techniques for the modulation of gene expression (shRNA and siRNA) were used to confirm the results of observational studies. Immunoprecipitation, in-gel or in-solution digestions and HLA-associated peptide purification were all performed in primary DCs isolated from healthy donors and analysed on an ultra-high performance liquid chromatography system. Cellular analysis of cross-presentation experiments was performed using CD8^+^ T cells from OT-1 C57BL/6 TCR-transgenic mice or human HLA-A2 NY-ESO-1_157-165_ CD8^+^ T cell clones and analysed by flow cytometry. The sample size is outlined in the figure legends.

### Generation of human monocyte-derived dendiritic cells and cell lines

Human monocytes were purified from peripheral blood mononuclear cells (PBMCs) from healthy donors by positive immunoselection with anti-CD14-conjugated MACS beads (Miltenyi Biotec). Monocytes were also purified from PBMCs from either HLA-A_2_ WT *NOD2* donors or HLA-A_2_ homozygous mutant *NOD2* Crohn’s patients. NIHR IBD Bioresource selected HLA-A_2_ Crohn’s patients expressing specific *NOD2* polymorphisms. Samples were collected in Oxford University NHS Foundation trust following written informed consent. Ethical approvals: (REC reference:16/YH/0247) and (REC reference: 09/H1204/30). Dendritic cells (DCs) were generated by culturing monocytes for 5 days with IL-4 and GM-CSF (Peprotech). Immature DCs were harvested on day 5 of culture. Bone marrow derived dendritic cells (BMDCs) were extracted from tibia and fibula bones of euthanized mice, passed through a 70-μm cell strainer, and cultured for 7 days in DC medium (RPMI 1640 medium with 10% FCS, kanamycin sulfate, MEM nonessential amino acids, sodium pyruvate, glutamine, 2-mercaptoethanol (55 mM) (all Life Technologies), and supplemented with recombinant mouse GM-CSF and IL-4 (20 ng/ml, Peprotech). 293/NOD2 cells were obtained by stable transfection of HEK293 cells with the pUNO-hNOD2 plasmid which expresses the human NOD2 gene (Invivogen). THP1 cells were obtained from ATCC.

### Cells and mice

C57BL/6 mice were from Envigo RMS Inc. (Bicester, Oxfordshire, UK). OT-1 C57BL/6 TCR-transgenic mice recognizing the H-2K^b^-restricted ovalbumin epitope SIINFEKL were from the Jackson Laboratory (Bar Harbor, Maine, USA). Mice were maintained at Biomedical Services, John Radcliffe Hospital. All animal studies were conducted with appropriate UK Home Office licenses and approval from the Oxford local ethics committee.

### Reagents and antibodies

PcDNA5 HA-PI31, pcDNA5 GFP-PI31 and pCMV-FLAG TBK1 were obtained from University of Dundee. SIINFEKL peptide, amino acids 257-264 within ovalbumin (OVA), and soluble OVA were obtanined from Sigma-Aldrich. Carboxyfluorescein diacetate succinimidyl ester CFSE was purchased from Biolegend. MDP, PAM_3_CSK_4_, LPS, R848, BX795 and Amlexanox were purchased from Invivogen. The antibodies used in the study include: rabbit anti-human PI31 (Atlas Antibodies, HPA041122), rabbit anti-mouse PI31 (Abcam, ab187200); rabbit anti-human P38 (9212), rabbit anti-mouse/human actin (12620), anti-human TBK1 (3504), anti-human IRF-3 (11904), rabbit anti-tubulin (5335), mouse anti-human PSMB8/LMP7 (13726), rabbit anti- human PA28α (9643), rabbit anti-human PSMD2 (25430), rabbit anti-human PSMA2 (11864), rabbit anti-human TANK (2141), rabbit anti-phospho-NF-κB p65 (3033), rabbit anti-human IκBα (9242), mouse anti-human phospho-IκBα (9246), rabbit anti-mouse ERp72 (5033), rabbit anti-mouse LAMP1 (3243), rabbit anti-FLAG (14793), rabbit anti-HA (3724), rabbit anti-GFP (2956), anti-mouse (7076) and anti-rabbit (7074) IgG HRP-linked secondary antibodies were all from Cell Signaling; rabbit-anti-human SEC16A (Abcam, ab70722), rabbit anti-mouse SEC16A (Holzel diagnostika, KIAA0310), mouse anti-human PSMA2 (Enzo, BML-PW8105-0100), mosue anti-human PSMA4 (Enzo, BML-PW8120). Anti-mouse CD3ε (Clone 145-2C11), anti-mouse CD8a (Clone 53-6.7), anti-mouse CD69 (Clone H1.2F3), anti-mouse MHC-II (Clone Af6-120.1), anti-mouse CD86 (GL1) and anti-mouse CD40 (Clone 3/23) were all from Biolegend.

### Cell stimulations, phosphoenrichment and immunoblots

Dendritic cells were left unstimulated or stimulated with 10 μg/ml MDP or 1 μg/ml PAM_3_CSK_4_ or both at the indicated time points. In some experiments, other PRR ligands were used including LPS 100 ng/ml, and R848 1 μg/ml (Invivogen) or cells were treated with the small molecule inhibitors Ponatinib (50nM) or BX795 (1mM) for 1 hr. Following stimulation, cells were harvested on ice and washed once with cold Hanks Buffered Saline (HBS). Phosphoenrichment was then performed using a cellular phosphoenrichment kit (Qiagen). Briefly, cells were lysed in phosphoprotein lysis buffer containing 0.25% CHAPS with phosphatase inhibitor cocktail 3 (Sigma), protease inhibitor tablet (Qiagen) and the nuclease 0.0002% Benzonase (Qiagen) at 4°C for 40 min. Samples were centrifuged and the supernatants harvested for protein quantification by BCA. Aliquots of whole cell lysate (WCL) were kept for subsequent immunoblots. Samples were then loaded onto the phosphoenrichment columns, before eluting the phosphoenriched fraction using an elution buffer containing 0.25% CHAPS (Qiagen). Following concentration of the eluted fraction using 9k molecular weight cut-off concentrator columns (Thermo Scientific), protein concentration was measured by BCA. LDS sample buffer (Life Technologies) and dithiothreitol (Sigma) was added, before heating to 70°C for 5 min. Protein samples were separated by SDS-PAGE and transferred onto PVDF or nitrocellulose membranes using iBlot2 dry blotting system (Invitrogen). Membranes were blocked in 5% (w/v) BSA or non-fat dry milk diluted in TBST for 1 hour at room temperature (RT). Primary antibodies were then added and incubated at manufacturers recommended dilution, temperature and time. Membranes were washed in TBST three times for 5-10 min. Species-specific HRP-conjugated secondary antibody was added to TBST with 5% (w/v) non-fat dry milk at the manufacturers recommended dilution. Membranes incubated in this solution for 1 hour at RT with gentle mixing. Membranes were washed 4 times for 5-10 min in TBST and developed by enhanced chemiluminescence (ECL) solution (Amersham). Densitometric analysis of Western blots was performed using ImageJ software.

### ShRNA lentiviral transduction and siRNA transfection

Short hairpin RNA lentiviral particles were produced and transduced following the RNAi Consortium (TRC) protocols. ShRNA containing pLKO.1 vectors targeting *NOD2* (SHCLND-NM_022162), *PI31* (SHCLND-NM_006814), *TBK1* (SHCLND-NM_013254) or non-Target shRNA Control Plasmid DNA were all obtained from Sigma (MISSION shRNA Plasmid DNA). Briefly, HEK293T packaging cells growing in 6 cm well plate were transfected with a mix of 1 μg packaging vector (psPAX2), 0.4 μg envelope vector (pMD2.G) and 1.6 μg hairpin-pLKO.1 vector (SHC016 control or gene specific shRNA. Fugene-6 (Promega) was used as transfection reagent. Cell culture medium containing lentiviral particles (LVP) was collected 48 h later and passed through a 0.45 μm filter (Sartorius). Virus preparations were then concentrated by centrifugation at 30,000 rpm for 90 min. Viral particles were added to cultured THP-1 cells in R10 together with 8 μg/ml Polybrene (Sigma) to improve transfection efficiency. Following incubation for 3 hrs at 37°C, the cells were harvested, washed, and resuspended at 1×10^6^ cells/ml in R10 media with antibiotics including puromycin (as selective antibiotic). After 10 days of continuous selection with puromycin, knockdown efficiency was assessed by immunoblot.

Transfection of human dendritic cells was performed by electroporation of SMARTpool ON-TARGETplus human Psmf1 (PI31) or non-targeting siRNAs (Dharmacon). Cells were resuspended in the solution provided with the kit (Invitrogen) followed by electroporation with Neon System kit (Invitrogen) using the following parameters: 1475 V, 20 ms, 2 pulses. BMDCs transfection was performed by electroporation of SMARTpool ON-TARGETplus mouse Psmf1, SEC16A or non-targeting siRNAs (Dharmacon) following the manufacturer’s instructions (Amaxa). Briefly, BMDCs were harvested at day 7 and resuspended in the electroporation solution provided with the kit. Cells were distributed per cuvette and electroporated. After 48 hrs, cells were harvested and knockdown analyzed by Western Blot.

### Immunoprecipitation, in-gel and in-solution digestions

Cells were lysed in IP lysis buffer (50 mM Tris-HCl (pH 8.0), 150 mM NaCl, 5 mM EDTA, 1% TritonX-100) containing protease and phosphatase inhibitor cocktail (Sigma). The lysate was centrifuged for 20 min at 4°C. The supernatant were collected and incubated with indicated antibodies or control isotype IgG for 4 hrs at 4°C. Protein G coupled magnetic beads (Invitrogen) were added and incubated for 1 hour at 4°C. The beads were washed with lysis buffer four times and protein samples eluted by incubation in LDS sample buffer with 50 mM DTT for 10 min at 70°C and analysed by SDS-PAGE and Western blotting using indicated antibodies. Alternatively, Pierce IgG elution buffer (Pierce; 210044) was added to beads for 30 min at 4°C with gentle agitation. For in-gel digestion, gel bands were cut into 1-2 mm^3^ cubes and were destained overnight with 50% methanol and 5% acetic acid in water. The samples were dried with acetonitrile and successively reduced with 10 mM DTT and alkylated with 50 mM iodoacetamide for 30 min at room temperature. The cubes were washed with 100 mM ammonium bicarbonate, dried with acetonitrile and digested with 400 ng elastase in 25 mM ammonium bicarbonate overnight at 37°C. Peptides were then extracted using three consecutive incubations of 10 min with a 25 mM ammonium bicarbonate followed by 50% acetonitrile and 5% formic acid in water and then 85 % acetonitrile and 5% formic acid in water. The samples were dried and resuspended in 1% acetonitrile and 0.1% TFA in water for LC-MS/MS analysis. For in-solution digestion, 35 μl of each IP eluate were suspended in 175 μl of water, reduced with 5 mM dithiothreitol followed by alkylation with 20 mM iodoacetamide for 1 hour at room temperature. Protein was precipitated using chloroform-methanol and re-suspended in 6 M urea in 0.1 M Tris pH 7.8. Protein material was digested with 600 ng Trypsin (Promega) overnight at 37°C at 300 rpm. The samples were desalted using C18 cartridge (Waters). Briefly, samples were conditioned with buffer A (1% acetonitrile, 0.1% TFA in water) prior equilibration with buffer B (65% acetonitrile, 0.1% TFA in water). Acidified peptides were loaded onto the column, washed with buffer A and eluted with buffer B. The solution containing the peptides was dried and resuspended in 1% acetonitrile, 0.1% TFA in water for LC-MS/MS analysis.

### qRT-PCR

Total RNA was purified using the RNeasy Mini Kit (Qiagen). cDNA was synthesised using a High Capacity RNA-to-cDNA kit (Applied Biosystems). Both procedures were performed according to the manufacturers’ protocols. qRT-PCR was performed using Taqman chemistry (Applied Biosystems). Taqman probes for NOD2 (Hs01550753_m1) and RPLP0 (Hs99999902_m1) and TaqMan Gene Expression Master mix were used. Real-time PCR was performed using a Bio-Rad C1000 Thermal cycler CFX Realtime system (Bio-Rad). Relative NOD2 gene expression was calculated in comparison to the RPLP0 control.

### Kinase assay

The TBK1 kinase assay was performed using the TBK1 Kinase Enzyme System (Promega) and ADP-Glo Kinase Assay kit (Promega) using the following peptides as substrate: IHEQWEKANVSSPHREFPPA (wild type PI31), IHEQWEKANVASPHREFPPA (mutated in Alanin 152), IHEQWEKANVSAPHREFPPA (mutated in Alanin 153) and IHEQWEKANVAAPHREFPPA (mutated in Alanins 152 and 153). Kinase Detection reagent was added to convert ADP to ATP, and the newly synthesized ATP was converted to light using the luciferase/luciferin reaction.

#### HLA-I-associated peptide purification

All steps were carried out below 4°C. Briefly, cell pellets were lysed using 10 ml lysis buffer (1% Igepal 630, 300 mM NaCl, 100 mM Tris pH 8.0) per 10^9^ cells and homogenized by mild sonication. Lysates were cleared in two subsequent centrifugation steps, one at 300 *g* for 10 min to remove nuclei and the other at 15,000 *g* for 30 min to pellet other insoluble material. HLA complexes were captured using 1 ml W6/32-conjugated immunoresin (1 mg/ml) prepared in a column format at a flow rate of 1.5 ml/min and washed using subsequent runs of 50 mM Tris buffer, pH 8.0 containing first 150 mM NaCl, then 400 mM NaCl and no salt. HLA-peptide complexes were eluted using 5 ml 10% acetic acid and dried. Samples were loaded onto on a 4.6 × 50 mm ProSwiftTM RP-1S column (ThermoFisher) in 120 μl buffer A (0.1% formic acid in water) and eluted using a 500 μl/min flow rate over 10 min from 2 % buffer A to 35 % buffer B (0.1% formic acid in acetonitrile) with an Ultimate 3000 HPLC system (Thermo Scientific). One ml fractions were collected from 2 to 15 min. Protein detection was performed at 280 nm absorbance. Fractions up to 12 min that did not contain ß_2_-microglobulin were combined, dried and resuspended in 1% acetonitril, 1% formic acid in water for LC-MS/MS analysis.

#### Liquid chromatography tandem mass spectrometry (LC-MS-MS)

LC-MS/MS analysis was performed on an ultra-high performance liquid chromatography system (Dionex Ultimate 3000, Thermo Scientific) supplemented with a 75 μm x 50 cm PepMap column coupled to a Fusion Lumos (Thermo Scientific) mass spectrometer with an EASY-Spray source. For in-gel and in-solution samples, peptides were analysed using 1 hr gradient from 2-35 % can and 0.1% formic acid in water at a 250 nl/min flow rate. Collision induced dissociation (CID) was induced on most intense ions in top speed mode with a 3 s cycle time and a collision energy of 35. Ions were excluded from repeated isolation for 60 seconds. HLA-I-associated peptide preparations were analysed using a 1 hr linear gradient from 3-25% ACN, 0.1% formic acid in water at a flowrate of 250 nl/min. Higher-energy C-trap dissociation (HCD) was induced on ions with 2-4 positive charges in top speed mode with a 2 s cycle time and a stepped collision energy of 28±5. Singly charged ions were selected at lower priority and fragmented using a stepped collision energy of 32±5. Ions were excluded from repeated fragmentation for 30 seconds.

#### MS data analysis

MS data was analysed using Mascot 2.5 and Peaks 7.5. Search parameters were set as follows: in-solution IP samples - enzyme specificity: trypsin, fixed modification - carbamidomethylation, precursor mass tolerance - 5 ppm, fragment mass tolerance - 0.3 Da. Data acquired from elastase digested material was searched with identical parameters using no enzyme specificity. A final FDR cut-off of 1% was applied to both datasets. Data generated from HLA-I-associated peptide preparations was searched using no enzyme specificity, no modifications and precursor mass tolerance of 5 ppm and fragment mass tolerance of 0.03 Da. All data was searched against all human protein sequences in the Uniprot database. FDR was <5% at a score cut-off of 15 for all samples. Quantitative analysis of the IP datasets was performed with Progenesis QI 3.0 (Nonlinear Dynamics). Protein quantitation was performed with non-conflicting peptides only.

### Purification of proteasomal proteins

Cells were lysed in proteasome lysis buffer (50 mM Tris-HCl (pH 8.0), 1 mM DTT, 5 mM EDTA, 5 mM MgCl_2_, 10% glycerol, 2 mM ATP) containing a protease and phosphatase inhibitor cocktail by five freeze and thaw cycles. Lysates were sonicated and centrifuged for 20 min at 4ºC. The supernatants were collected and pre-cleared by agarose beads for 1 hr at 4 ºC. 900 μg of protein was incubated with anti-PSMA2 (MCP21) antibody (BML-PW8105) or control mouse IgG (sc-2027) overnight at 4 ºC. Then, protein A/G PLUS-Agarose (sc-2003) was added and rotated for 2 hrs at 4 ºC. The beads were washed four times by proteasome lysis buffer with 0.1% NP-40. Co-immunoprecipitated proteins were eluted by incubation in LDS sample buffer with 50 mM DTT for 10 min at 70 °C. Subsequently, eluted proteins were analysed by SDS-PAGE and Western blotting.

### Confocal Microscopy

Following electroporation of pcDNA5 GFP-PI31 (Amaxa), cells were placed on poly-L-lysine-coated glass coverslips and incubated at 37°C in an atmosphere of 5% CO2. Cells were left unstimulated or stimulated with 10 μg/ml MDP and 1 μg/ml Pam_3_CSK_4_ for 30 min. After extensive washing with cold PBS, cells were fixed with 2% (vol/vol) paraformaldehyde during 10 min at 4°C and quenched by adding 0.1 M glycine. Cells were permeabilized in PBS containing 0.05% (vol/vol) saponin and 0.2% (vol/vol) BSA for 20 min at room temperature, washed, and incubated first with primary antibodies for 1 hr and then with secondary antibodies for 45 min at RT. After washing, the coverslips were mounted with Vectashield (Vector Laboratories). Image acquisition was performed on a Zeiss inverted confocal LSM 880 microscope. Analysis was performed using Fiji software.

### Phagosome isolation and analysis

BMDC were pulsed with equal numbers of unconjugated 3 um magnetic streptavidin microspheres or magnetic streptavidin microspheres (Bangs Laboratories) conjugated with biotinylated MDP (10 ug/ml) and PAM_3_CSK_4_ (1 ug/ml) or biotinylated LPS (100 ng/ml) in a 1:2 ratio of BMDC: beads for 3 hrs. Cells were suspended in homogenization buffer (HB) (250 mM sucrose, 0.5 mM EGTA, and 20 mM HEPES/KOH) containing protease and phosphatase inhibitor cocktail and disrupted by sonication. Phagosomes containing the magnetic beads were isolated from homogenate using a magnet (Invitrogen). For western blot analysis, the magnetic beads were lysed in elution buffer (50 mM Tris, pH 7.9, 300 mM NaCl, 1% Triton) along with protease and phosphatase inhibitor cocktail for 30 min at 4ºC with gentle agitation. Protein concentrations were measured by Bradford assay. Isolated phagosome proteins (2 ug) and whole cell lysate (15 ug) were analysed by western blot.

### *In Vitro* Cross-Presentation Assays

BMDCs were left unstimulated or stimulated with 10 μg/ml MDP and 1 μg/ml PAM_3_CSK_4_ and incubated with sOVA or SIINFEKL peptide for 3 hrs. After extensive washing, BMDCs were incubated with CD8^+^ lymphocytes (72 hrs) isolated from C57BL/6 OT-1 transgenic mice and purified by negative immunomagnetic bead selection (Miltenyi-Biotec). Cross-presentation was evaluated by detecting expression of CD69^+^ on CD8^+^ lymphocytes by FACS.

### *In Vivo* inhibition of TBK1 activity and cross-presentation assay

Mice were injected intraperitoneally with Amlexanox (10mg/Kg, Invivogen). After 24 hrs, the same amount of Amlexanox was injected again intraperitoneally with 100 μg MDP (Invivogen) and 10ug of PAM_3_CSK_4_. DMSO diluted in endotoxin-free PBS was injected as a control. After 12 hrs, sOVA was injected intravenously (250 ug), and spleens were harvested after 3 hrs. DCs were purified by CD11c negative selection (Miltenyi). DC activation was determined by upregulation of the co-stimulatory molecules MHC-II, CD86 and CD40. DCs were co-cultured with purified CFSE-OT-I CD8^+^ T cells for 3 days. T cell proliferation was measured by CSFE staining by FACS.

### Generation of HLA-A2 NY-ESO-1_157-165_ T cell clones

NY-ESO-1 specific CD8^+^ CTL clone was sorted directly using HLA-A2 NY-ESO-1_157-165_ tetramers from a melanoma patient, as previoulsy described (28). After expansion, CTLs were stimulated with a mixture of allogeneic irradiated PBMCs and LG2 cells in the presence of 5 μg/ml phytohaemagglutinin and cultured over 16 days at 37°C, 5% CO2 in RPMI-1640 (Invitrogen) supplemented with 10% (v/v) human serum, 2 mM Glutamax, 50-100 U/ml penicillin and 100 μg/ml streptomycin, 10 mM HEPES, 50 μΜ 2-mercaptoethanol, 1 mM sodium pyruvate, non-essential amino acids and 400 U/ml recombinant human IL-2 (Novartis).

#### Statistical and data analysis

Data were analyzed with Graph Pad Prism, and the Student’s *t* test or one-way ANOVA for multiple comparisons was used to compare data sets throughout this study. *P* < 0.05 was considered significant.

## Results

### NOD2 and TLR2 stimulation induce PI31 phosphorylation

We first delineated NOD2 and TLR2 signaling in primary human DCs using quantitative phospho-proteomic analysis. We compared NOD2 stimulation with TLR2 stimulation alone or in combination. After stimulating cells with either MDP, PAM_3_CSK_4_ or a combination of both ligands, we lysed the cells and phosphoenriched (PE) the lysates. Phospho-enriched lysates were then subject to quantitative phosphoproteomic analysis by liquid chromatography-tandem mass spectrometry (LC-MS/MS) (fig. 1A). Using computational analysis, we identified proteins as differentially abundant, if the median among 5 donors was either 1.5-fold up- or downregulated compared to unstimulated controls. Among these differentially regulated proteins, we identified 134 proteins when stimulated with MDP (38 up-regulated, 96 down-regulated), 123 proteins when stimulated with PAM_3_CSK_4_ (19 up-regulated, 104 down- regulated) and 164 proteins when stimulated with both MDP and PAM_3_CSK_4_ (48 up-regulated, 116 down-regulated) (fig.1B; fig. S1). PI31 was highly abundant among the proteins differentially regulated in response to MDP (Table 1, fig.1C).

**Table 1.**
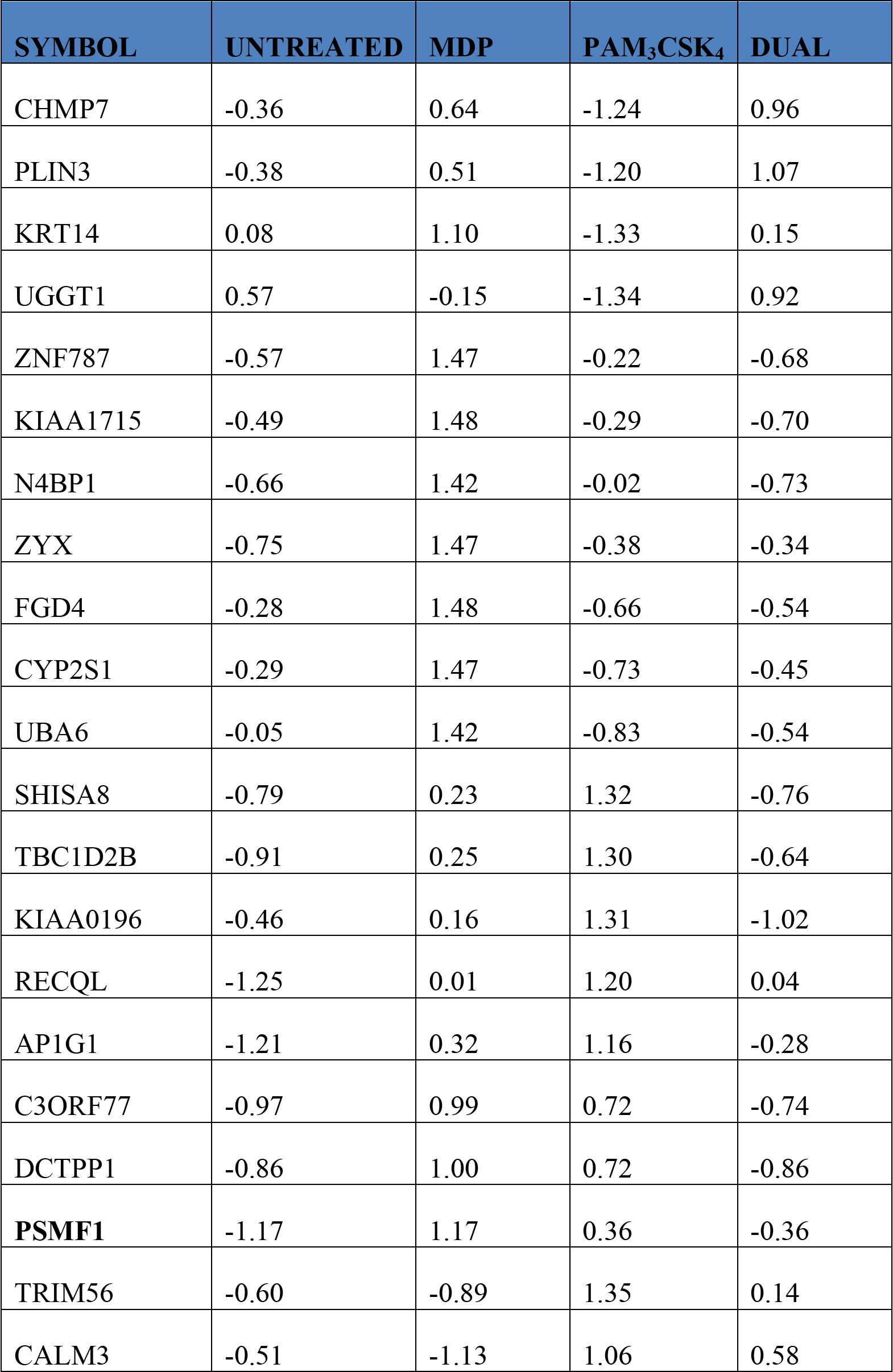
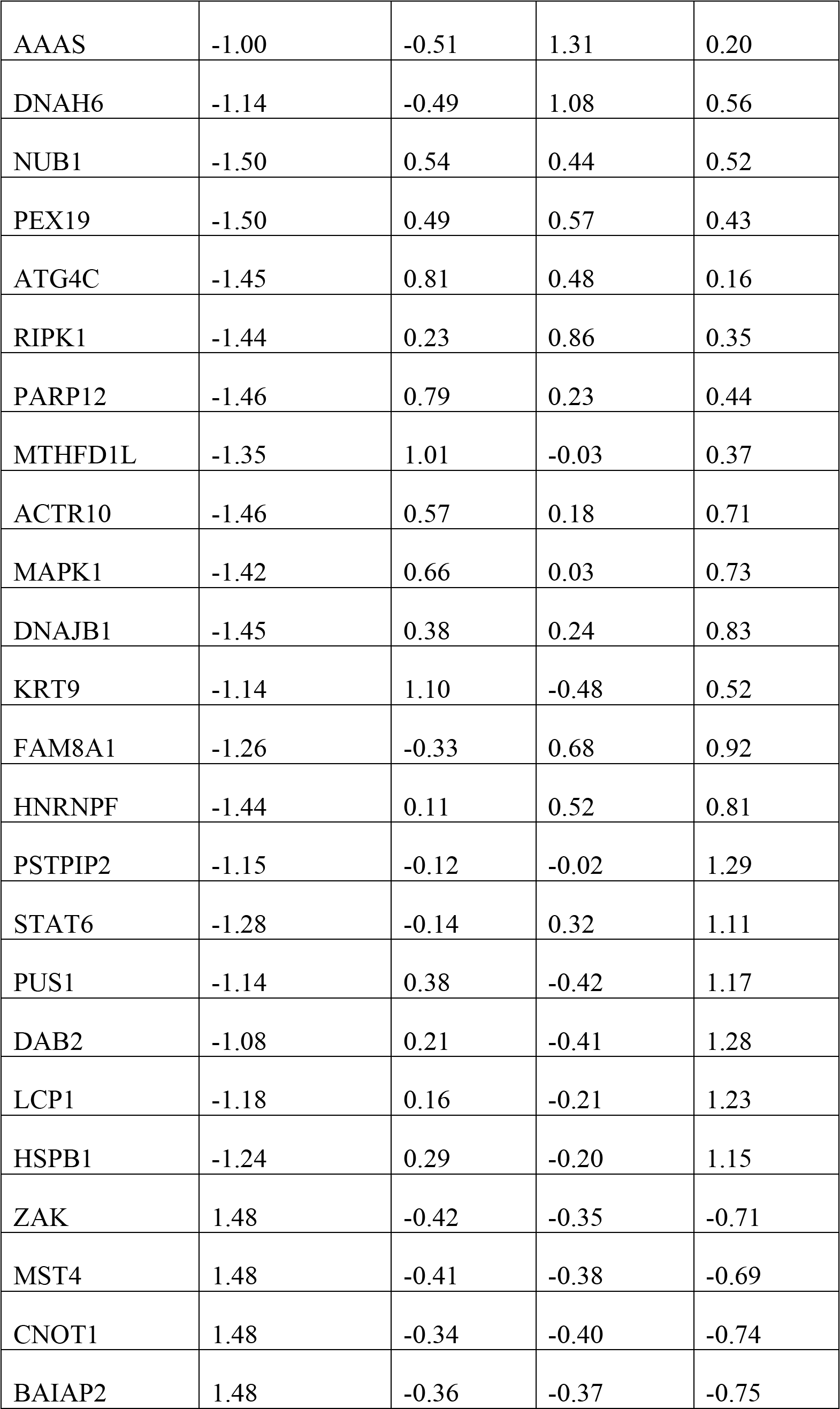
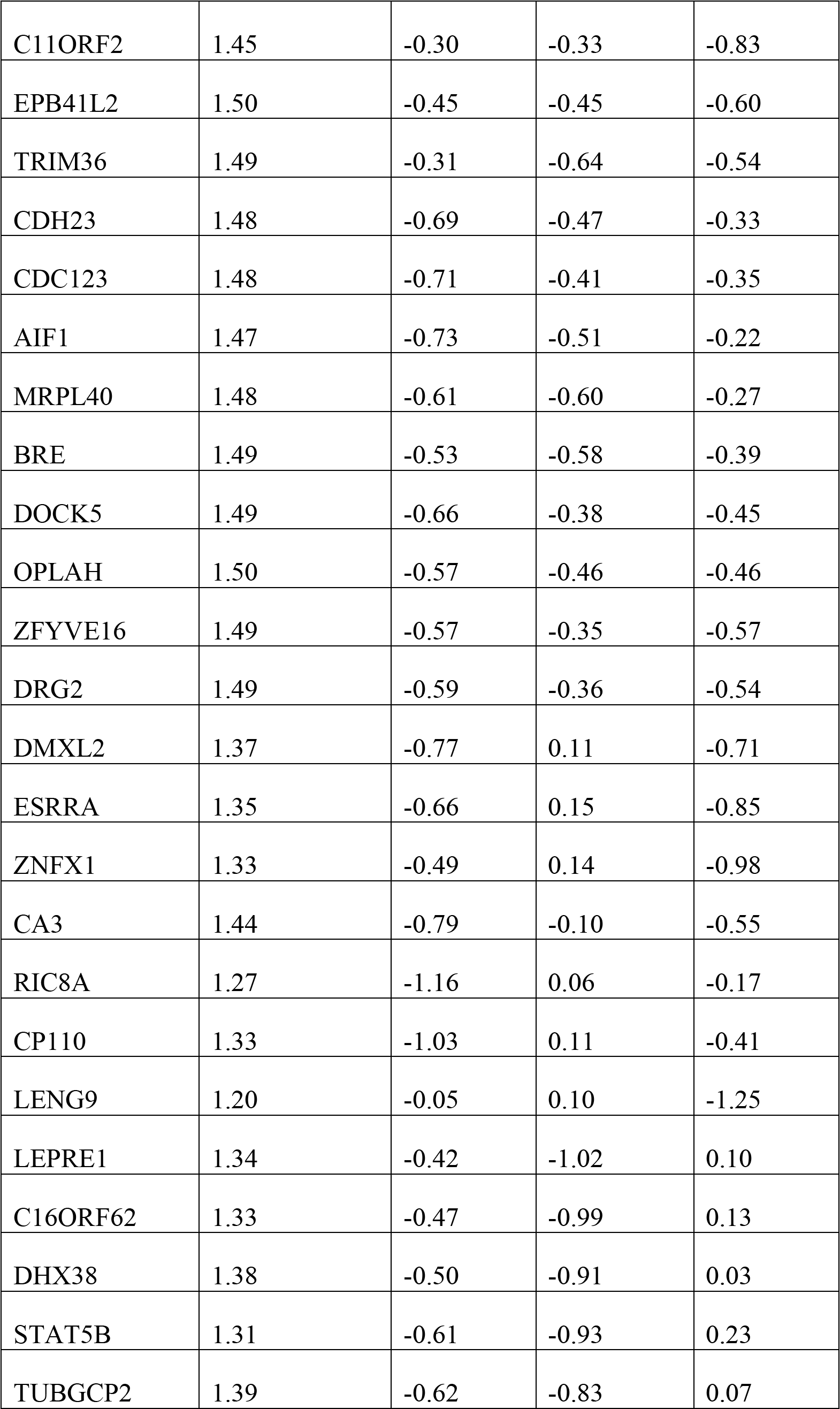
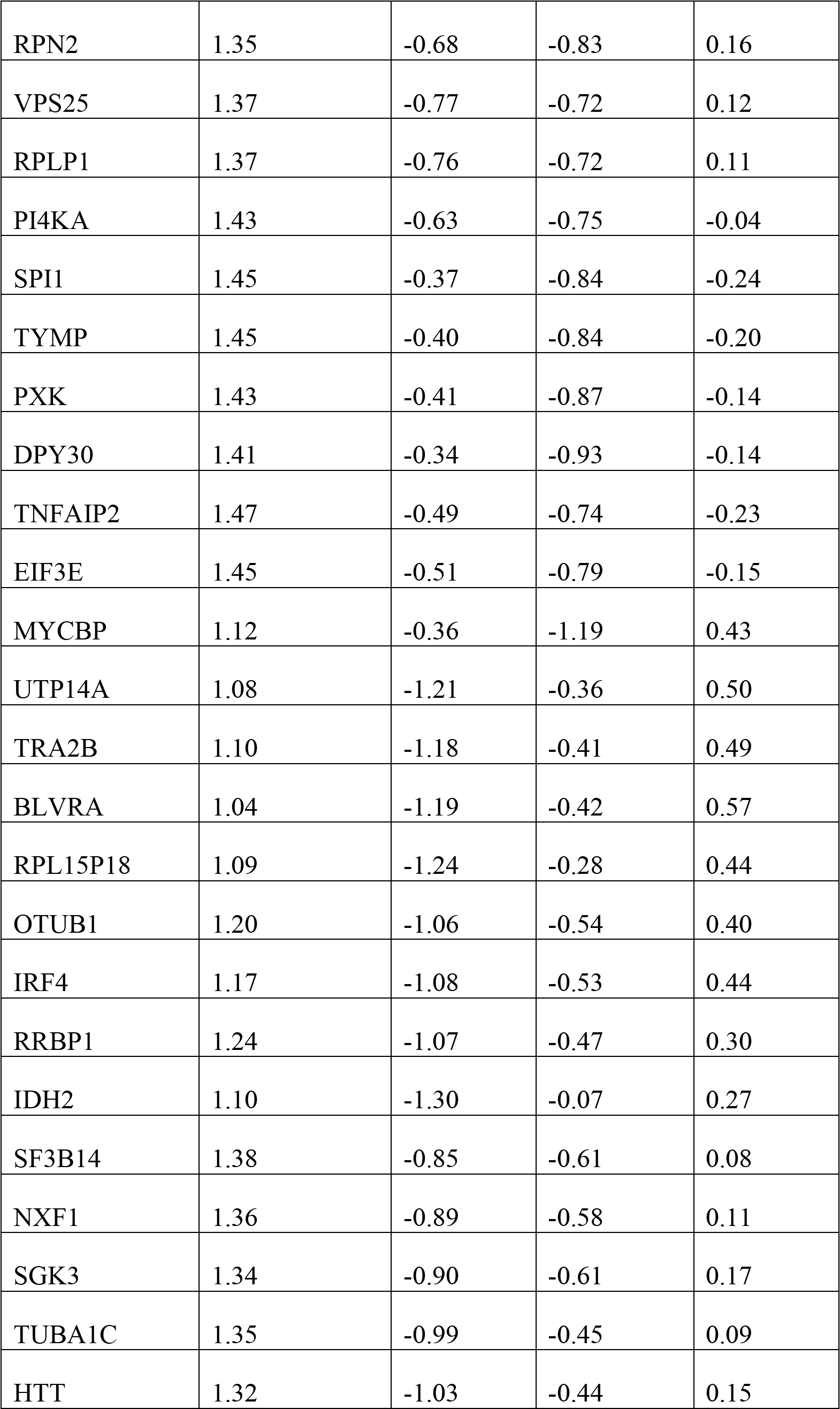
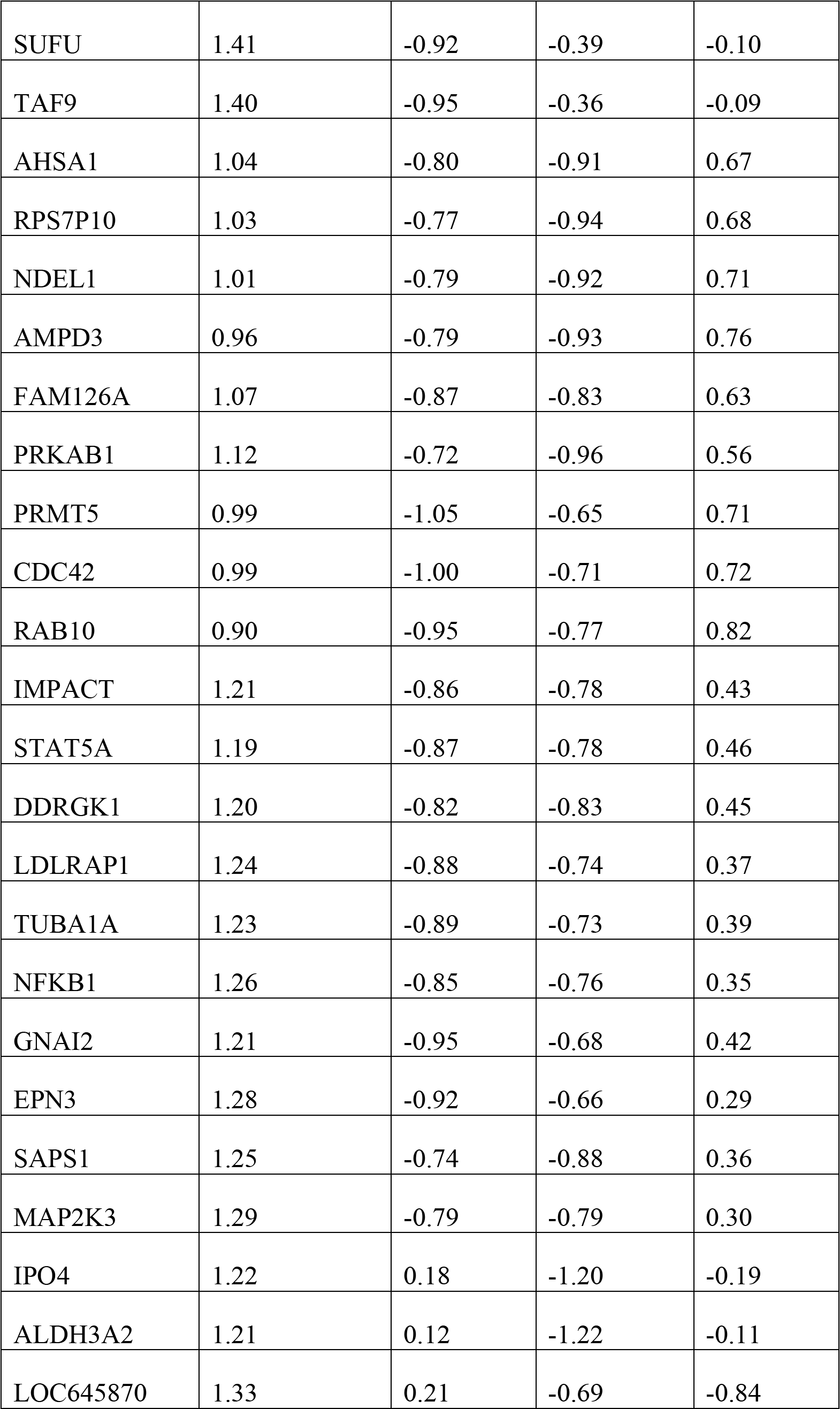
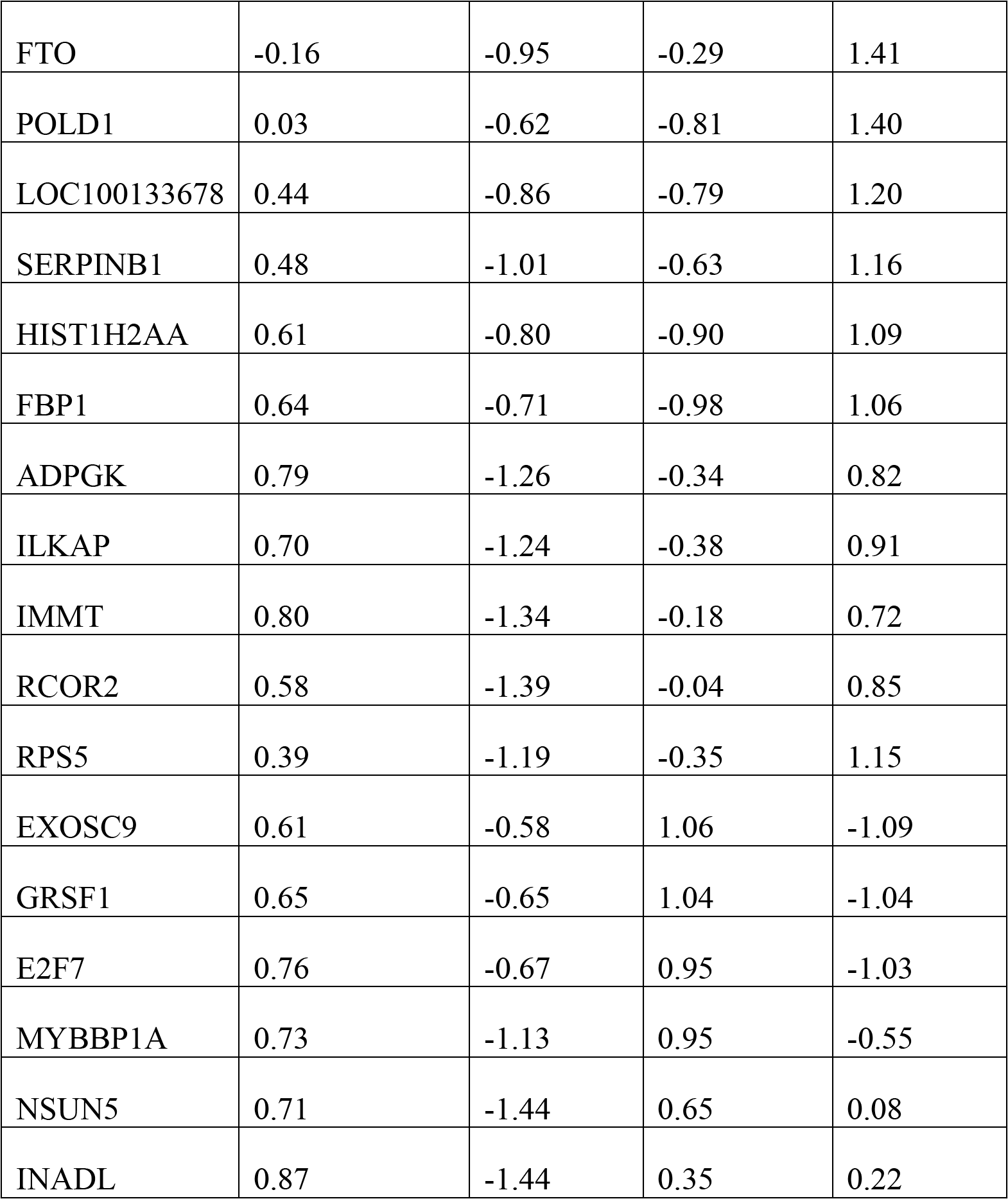
Table showing the z-score normalized abundances of hierarchical clustering presented in fig.1b (main text)

**Figure 1.**
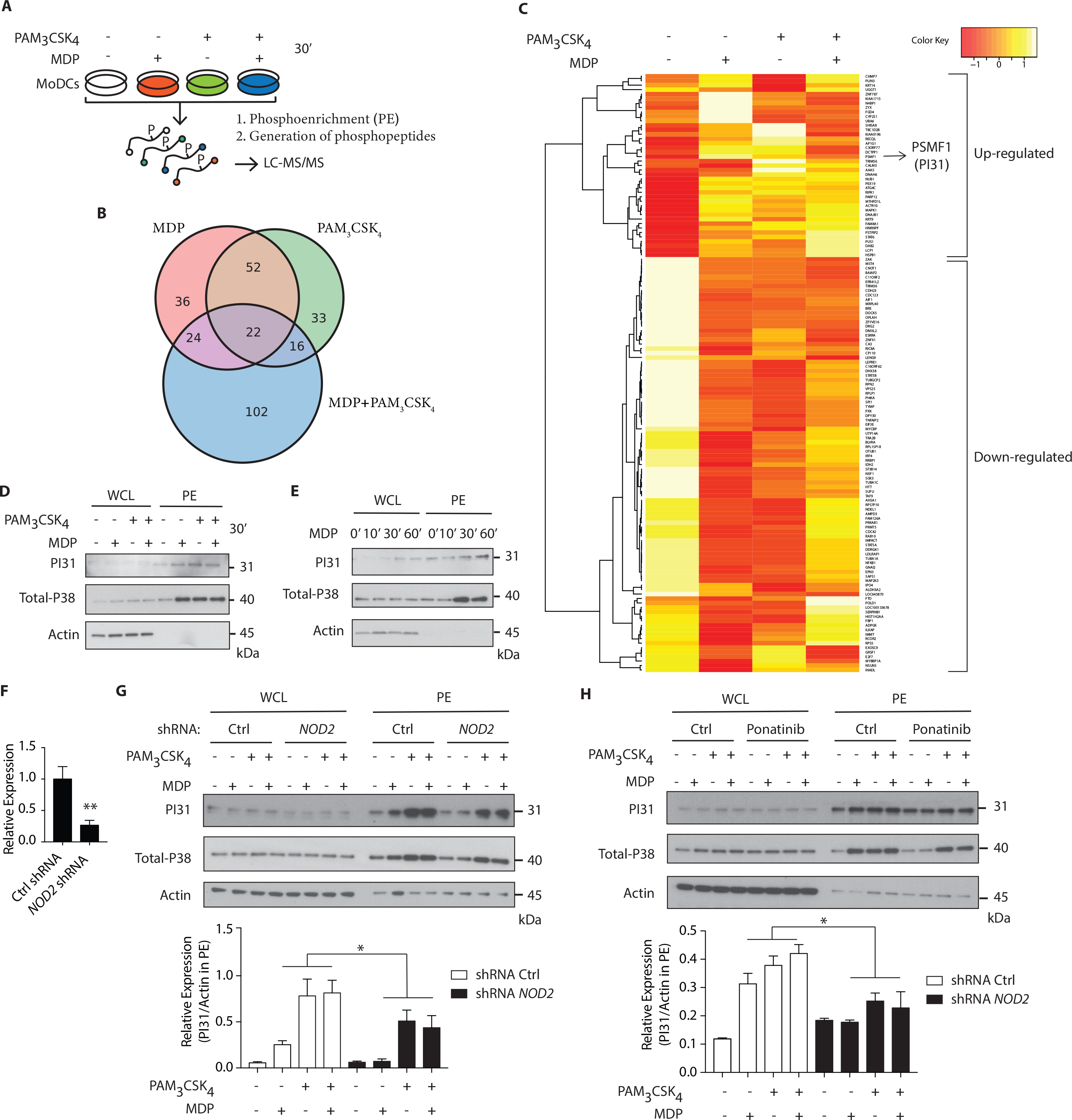
Quantitative phosphoproteomic analysis reveals PI31 phosphorylation on NOD2 and TLR2 stimulation in human DCs. (**A**) Schematic demonstrating the strategy for performing quantitative phospho-proteomics in DCs stimulated for 30 min with MDP (10 μg/ml), PAM_3_CSK_4_ (1 μg/ml) and MDP + PAM_3_CSK_4_ combined, followed by phosphoenrichment (PE) and generation of phosphopeptides for LC-MS/MS. (**B**) Venn diagram of differentially regulated proteins (up- or down-regulated) following MDP, PAM_3_CSK_4_ and MDP + PAM_3_CSK_4_ stimulation. (**C**) Hierarchical clustering showing proteins with median phosphopeptides either 1.5-fold up- or down-regulated in response to MDP compared to unstimulated Ctrl and selected as differentially abundant (n=5). (**D-E**) Immunoblot analysis of PI31 and P38 (positive control) in whole cell lysates (WCL) and PE following stimulation of DCs at indicated time points by NOD2 and TLR2 ligands alone or in combination as indicated: MDP (10 μg/ml), PAM_3_CSK_4_ (1 μg/ml). Immunoblot data are from one experiment representative of three separate experiments. Experiments using primary DCs were performed using two separate donors each time. (**F**) THP1 cells were transduced with control or *NOD2*-targeting lentiviral shRNAs and analysed for *NOD2* expression by qPCR analysis. Data represent the mean ± s.e.m. (*n* = 3); \*\**P*<0.001 Student’s *t* test. (**G**) Immunoblots analysis of PI31 and P38 in whole cell lysates (WCL) and PE of THP1 transduced with control or *NOD2* shRNA and stimulated with MDP (10 μg/ml), PAM_3_CSK_4_ (1 μg/ml), and MDP + PAM_3_CSK_4_ for 30 min. Immunoblots data are from one experiment representative of three separate experiments. Densitometric analysis of PI31 in PE band intensity normalized with actin (*n*=3). Data represent the mean ± s.e.m. (*n* = 3); \**P*<0.05 Student’s *t* test. (**H**) Immunoblots analysis of PI31 and P38 in whole cell lysates (WCL) and PE from DCs treated with Ponatinib (50nM) for 1 hr following stimulation with MDP (10 μg/ml), PAM_3_CSK_4_ (1 μg/ml), and MDP + PAM_3_CSK_4_ for 30 min. Densitometric analysis of PI31 in PE band intensity normalized with actin (*n*=3). Data represent the mean ± s.e.m. (*n* = 3); \**P*<0.05 Student’s *t* test.

PI31 has been linked to proteasome function (29,30) but no previous studies have reported its phosphorylation and association with innate immune pathways. Therefore, we speculated that PI31, given its link to proteasome, might be a suitable candidate to trigger MHC class I antigen presentation on NOD2 and TLR2 sensing. First, we confirmed that NOD2 activation by MDP, TLR2 activation by PAM_3_CSK_4_ or their combination, induced PI31 phosphorylation (fig. 1 D, E). We used phosphorylated MAPK (p38) in PE lysates as a positive marker of MAPK activation. We then determined whether NOD2 was required for PI31 phosphorylation. Using short hairpin RNAs (shRNA) targeting *NOD2*, we downregulated *NOD2* expression in THP-1 cells (fig. 1F). We assayed protein levels of phosphorylated PI31 following stimulation with MDP, PAM_3_CSK_4_ or a combination of both ligands. We found reduced PI31 phosphorylation after MDP exposure in NOD2 knockdown cells (fig. 1G). Reduced levels of PI31 phosphorylation were also observed following stimulation with PAM_3_CSK_4_ or a combination of both MDP and PAM_3_CSK_4_ suggesting that NOD2 may also amplify TLR2-dependent phosphorylation of PI31 (fig.1G).

NOD2 signaling requires RIPK2, but not the TLR adaptor MyD88 (31). So, we determined whether PI31 required RIPK2 for phosphorylation via NOD2. We pre-treated DCs with Ponatinib, a potent RIPK2 inhibitor (32,33), followed by stimulation with MDP, PAM_3_CSK_4_ or both for 30 min. We found decreased PI31 phosphorylation in cells treated with Ponatinib compared to untreated cells, which demonstrates that NOD2 mediated PI31 phosphorylation requires RIPK2 (fig. 1H).

Formation of the NOD2-RIPK-2 complex following MDP recognition induces ubiquitination of IκB kinase-γ, which then recruits TAK-1 and activates the downstream targets, NF-κB and MAPK (34). Next, we determined whether engaging PI31 through NOD2 and TLR2 co-stimulation interfered with NF-κB and MAPK signalling. We assayed levels of phospho-IκBα, phospho-p65 and phospho-p38 after stimulation with MDP and PAM_3_CSK_4_. shRNA knockdown of PI31 in THP-1 cells did not modulate activation of either NF-κB or MAPK signaling (fig. S2A, B). We also examined whether PI31 modulates NOD2/TLR2-induced secretion of pro-inflammatory cytokines. PI31 expression was downregulated in primary DCs using siRNA prior to stimulation with MDP and PAM_3_CSK_4_ or LPS. PI31 knockdown did not modulate cytokine secretion as detected by levels of TNF and IL-6 (Fig. S2C, D). These data demonstrate that stimulation of DCs with NOD2 and TLR2 ligands induces phosphorylation of PI31, a protein that alters the function of the proteasome. PI31 activation by NOD2/TLR2 signaling did not affect NF-κB and MAPK activation or cytokine production in DCs.

### TBK1 mediates NOD2 and TLR2 mediated PI31 phosphorylation

NOD2 stimulation triggers the formation of a protein complex containing RIPK2 and TRAF3 (tumour-necrosis-factor-receptor-associated factor 3), an E3 ligase that mediates Lys63-linked ubiquitination, through a serine/threonine-protein kinase TBK1 dependent mechanism (35,36). To determine whether TBK1 forms a protein complex with PI31, we co-expressed HA-tagged PI31 and FLAG-tagged TBK1 in HEK293 cells expressing human NOD2 (hNOD2) and stimulated the cells with MDP. We immunoprecipitated HA-PI31 from cellular extracts and detected the presence of FLAG-TBK1 by immunoblot of the immunoprecipitated material. This result suggests that PI31 and TBK1 are in the same complex (fig. 1A) with increased detection of TBK1 co-immunoprecipitating with PI31 following MDP treatment. We then expressed FLAG-TBK1 and GFP-PI31 in hNOD2 HEK293 cells and stimulated them with MDP. Co-immunoprecipitation of FLAG-TBK1 and GFP-PI31 confirmed the presence of TBK1 and PI31 in the same complex. We found increased levels of GFP-PI31 following MDP stimulation, suggesting an increased recruitment of PI31 co-immunoprecipitating with TBK1 following NOD2 activation (fig. 2B).

**Figure 2.**
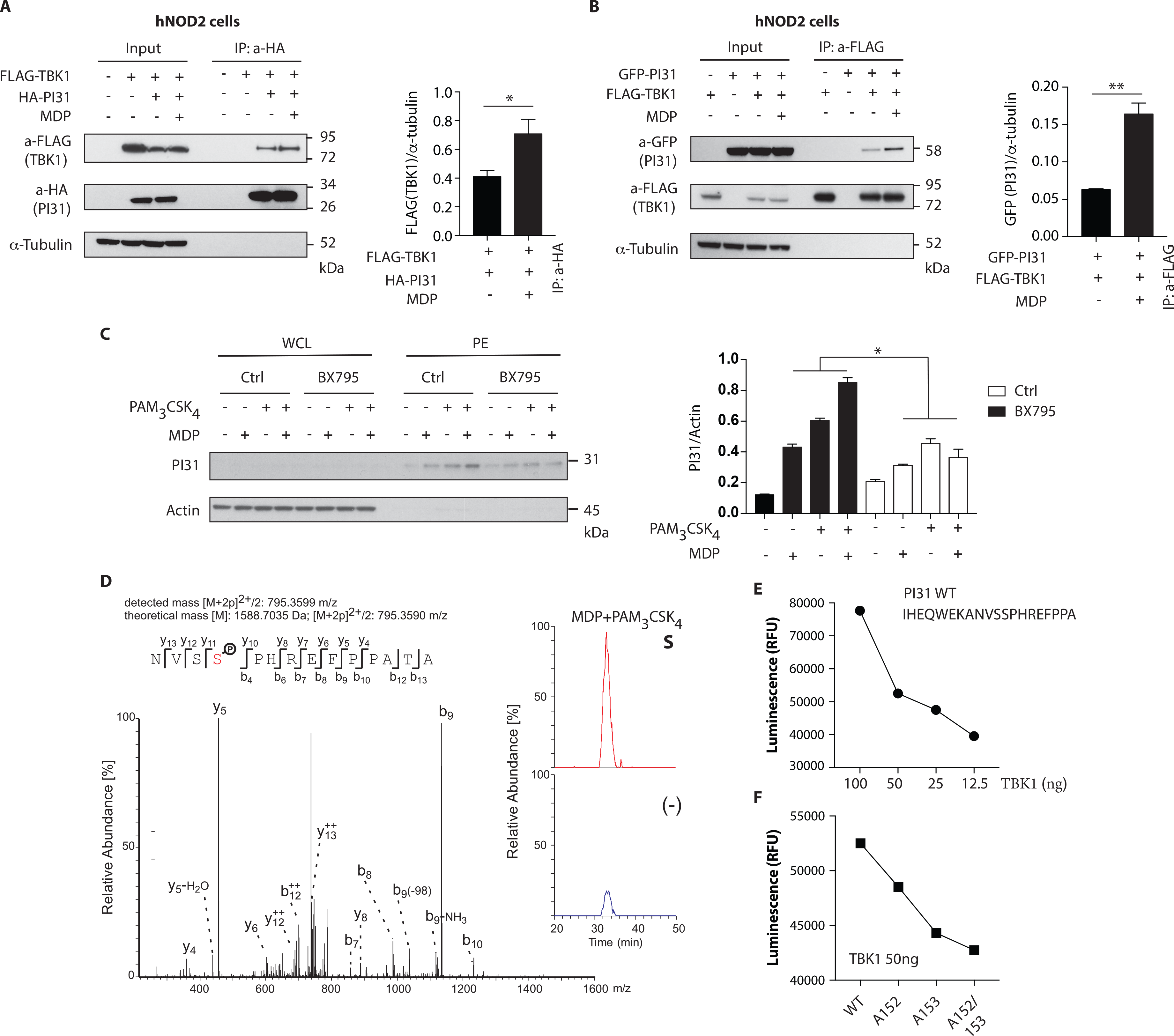
TBK1 mediates NOD2 and TLR2 mediated PI31 phosphorylation. (**A**) Constructs were transfected into HEK293 cells expressing human NOD2 (hNOD2) and cells stimulated with MDP for 30 min. Cell lysates were immunoprecipitated with anti-HA antibodies (PI31) and western blotted with anti-Flag (TBK1) antibodies. Densitometric analysis of Flag-TBK1 band intensity normalized with α-tubulin (*n*=3). Data represent the mean ± s.e.m. (*n* = 3); \**P*<0.05 Student’s *t* test. (**B**) Cell lysates were immunoprecipitated with anti-Flag antibodies (TBK1) and immunobloted with anti-GFP antibodies (PI31). Densitometric analysis of GFP-PI31 band intensity normalized with α-tubulin (*n*=3). Data represent the mean ± s.e.m. (*n* = 3); \*\**P*<0.01 Student’s *t* test‥ (**C**) Immunoblots of PI31 in whole cell lysates (WCL) or phosphoenriched fractions (PE) from treated primary DCs treated with BX795 (1 μM) for 1 hr following 30 min stimulation with MDP (10 μg/ml), PAM_3_CSK_4_ (1 μg/ml) or both. Immunoblots represent three independent experiments. Experiments using primary DCs were performed using two separate donors each time. Densitometric analysis of PI31 band intensity in PE normalized with actin (*n*=3). Data represent the mean ± s.e.m. (*n* = 3); \**P*<0.05 Student’s *t* test. (**D**) Immunoprecipitation of PI31 from THP1 stimulated with MDP/PAM_3_CSK_4_ was digested to in solution followed by LC-MS/MS analysis. Peptide 795.3599 m/z was identified as the NVSSPHREFPPATA phosphopeptide from PI31, where serine-153 was phosphorylated. Quantitation of unstimulated and MDP/PAM_3_CSK_4_ stimulated samples showed significant enrichment of the peptide in stimulated samples as shown by the percentage relative abundance. (**E**) Recombinant TBK1 enzyme was titrated with 50mM ATP and PI31 peptide (1mg/ml) prior to measuring kinase activity. (**F**) TBK1 kinase activity following incubation of recombinant TBK1 with wild type or mutated PI31 peptides. Data represent two independent experiments.

We next examined whether PI31 phosphorylation following NOD2 and TLR2 activation requires TBK1 kinase activity. We pre-treated primary human DCs with a TBK1 inhibitor, BX795, which specifically inhibits the catalytic activity of TBK1 (37), followed by DC stimulation with MDP, PAM_3_CSK_4_ or a combination of both ligands. Pharmacologic inhibition of TBK1 significantly reduced PI31 phosphorylation after NOD2 and TLR2 stimulation. (fig. 2C).

Next, we sought to identify the residue on which PI31 is phosphorylated following NOD2 and TLR2 activation. We immunoprecipitated endogenous PI31 from cellular extracts of THP1 cells stimulated with MDP and PAM_3_CSK_4_, and then conducted LC-MS/MS analysis on the immunoprecipitated material. Manual inspection of the mass spectrometry data identified the peptide 795.3599 m/z as the NVSSPHREFPPATA phosphopeptide from PI31 where serine 153 was phosphorylated. Quantitation of unstimulated and MDP/PAM_3_CSK_4_ stimulated samples showed significant enrichment of the peptide in stimulated samples indicated by the percentage relative abundance (fig. 2D). We next validated the mass spectrometry data by using a luminescent *in vitro* kinase assay that measures the amount of ADP formed from the TBK1 kinase reaction. First, we titrated the TBK1 enzyme in the presence of a PI31 sequence peptide containing both serines 153 and 152 (fig 2E). We examined mutated peptides for serine 152 (alanine 152), serine 153 (alanine 153) or both with the *in vitro* kinase assay, which confirmed PI31 was directly phosphorylated at serine 153 and, to a lesser extent, at serine 152 (fig 2F). These results demonstrate that TBK1 forms a complex with PI31 and may be required for its phosphorylation on serine 153 on NOD2 and TLR2 sensing.

### PI31 modulates the quantity of MHC class I loaded peptides in DCs

Proteasomes remove abnormal proteins from the cytosol and contribute to proteolysis required for MHC-class I antigen-processing (38-40). Most proteasomal degradation depends on recognition of poly-ubiquitinated proteins (41). As PI31 interacts with and inhibits proteasomes (30), we wanted to determine whether its phosphorylation via NOD2/TLR2 leads to its poly-ubiquitination and degradation by the proteasome. We stimulated DCs with MDP and PAM_3_CSK_4_ for 45 or 180 min and then collected the lysates for PI31 immunoprecipitation. Precipitates were immunoblotted using anti-ubiquitin (FK2) antibody. We did not detect poly-ubiquitination of PI31 in either non-stimulated or MDP/PAM_3_CSK_4_ stimulated samples (fig. S3A). To examine whether PI31 is degraded after its phosphorylation, we stimulated DCs and THP1 cells with MDP and PAM_3_CSK_4_ for 0, 0.5, 1, 24 and 48 hrs and assessed total levels of PI31 by immunoblot analysis. PI31 levels remained stable over time post-stimulation, which suggests it is not degraded by NOD2 or TLR2 sensing (fig. S3B, C).

PI31 can inhibit immunoproteasome formation and may maintain a balance between constitutive and immunoproteasome activity (29). So, we sought to determine whether PI31 expression in primary DCs would alter the generation of MHC class I bound peptides derived from proteasomal degradation. LC-MS/MS allows direct qualitative and quantitative evaluation of HLA class I-bound peptides from primary cells. We used siRNAs to knockdown PI31 in primary DCs, followed by capturing HLA class I complexes using W6/32-conjugated immunoresin (fig. 3A, B). Acid treatment abolished noncovalent interactions among the complex components. Using reverse-phase HPLC, we separated eluted peptides from the α-chain and β_2_-microglobulin of the HLA complexes and analysed eluted peptide fractions by LC-MS/MS. Mass spectrometry analysis showed that PI31 knockdown reduced the total number of peptides eluted compared to cells expressing PI31 (fig. 3C, D). The peptide motifs were not significantly modulated by PI31 suggesting that its effect is unlikely restricted to generating specific type HLA class I associated peptides (fig. 3E). We next tested whether PI31 modulates the number of assembled β_2_-microglobulin MHC-I complexes competent for surface antigen presentation. We used siRNAs to knockdown PI31 in primary DCs (fig. 3F), followed by measurement of β_2_-microglobulin before (lysate) and after (supernatants) capturing HLA class I complexes. The abundance of β_2_-microglobulin peptides detected by LC-MS/MS was not significantly modified by PI31 before or after capturing HLA-class I complexes (fig. 3G, H). This confirmed PI31 affects generation of different HLA class I associated peptides rather than quantity of surface MHC I.

**Figure 3.**
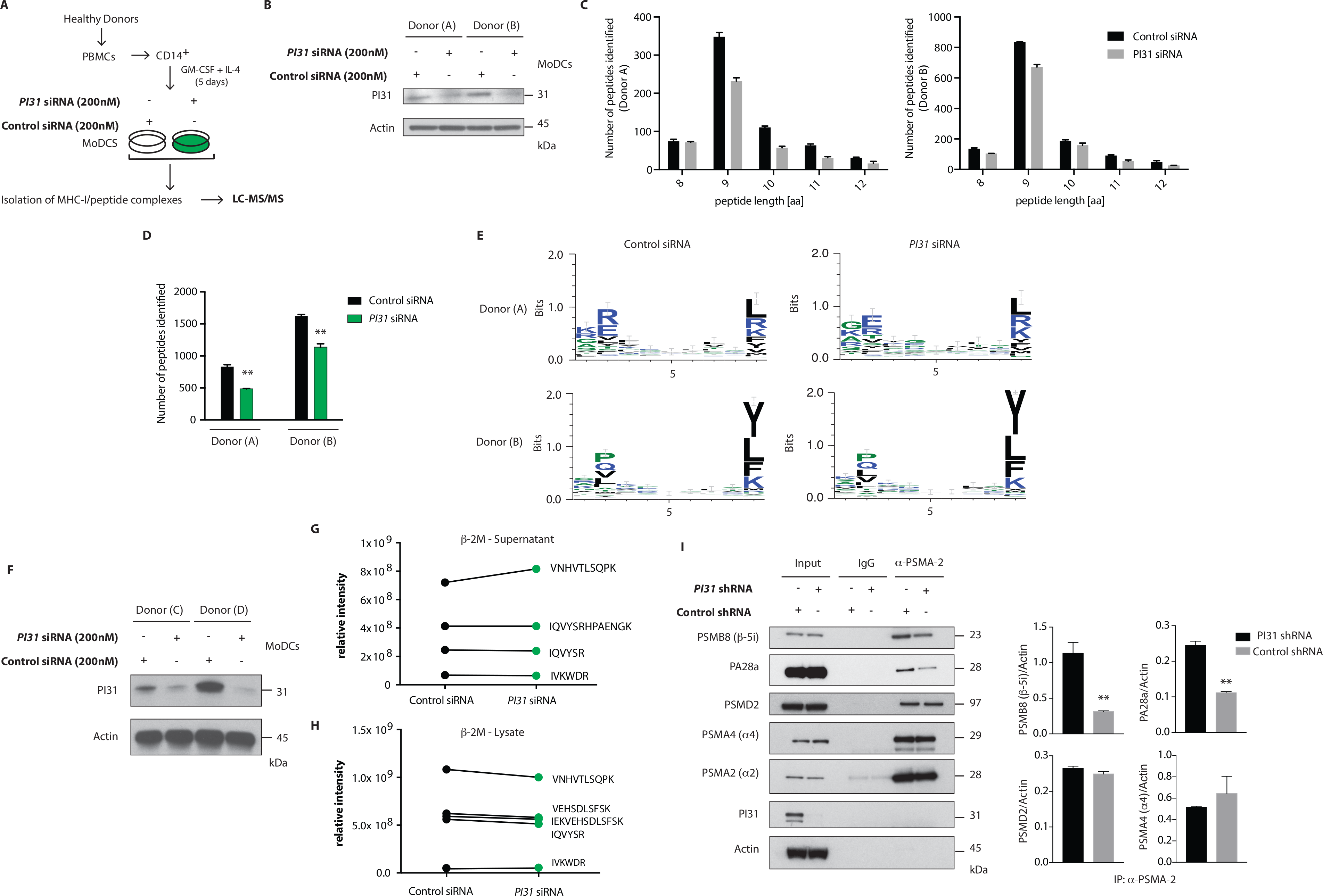
PI31 regulates the number of peptides loaded on MHC I and stabilizes the immunoproteasome. (**A**) Schematic of the strategy to assay peptides associated with MHC-I in Ctrl or PI31 knockdown DCs. (**B**) Immunoblot analysis of PI31 expression following transfection of primary DCs (donors A and B) with Ctrl non-targeting or *PI31* siRNAs (200nM). (**C**, **D**) Number of MHC-I-peptides identified in DCs from two healthy donors. Data represent mean ± s.e.m. (*n* = 2); \*\**P*<0.001 Student’s *t* test. Data represent an independent experiment using cells from two separate donors. (**E**) Effect of PI31 on HLA class I associated peptides. (**F**) Immunoblot analysis of PI31 expression following transfection of primary DCs (donors C and D) with Ctrl non-targeting or *PI31* siRNAs (200nM). (**G**, **H**) Relative intensities of β_2_-microglobulin peptides detected by LC-MS/MS before (lysate) and after (supernatants) capturing HLA class I complexes. (**I**) Cell lysates from THP1 cells transduced with Ctrl or *PI31* shRNAs immunoprecipitated using anti-PSMA-2 (MCP21) and immunocomplexes analyzed by western blot to determine the levels of PSMB8, PA28a, PSMD2, PSMA4, PSMA2 and PI31. Densitometric analysis of band intensities of PSMA-2 immunoprecipitation normalized with actin (*n*=3). Data represent the mean ± s.e.m. (*n* = 3); \*\**P*<0.01 Student’s *t* test.

Next, we investigated the mechanism through which PI31 modifies the number of HLA class I associated peptides by regulating constitutive and immunoproteasome subunit composition. Using THP-1 cells with knocked down PI31 by shRNA, we immunoprecipitated PSMA2 (MCP21), a 20S core alpha subunit, and examined the levels of constitutive or immunoproteasome subunits by immunoblotting. PI31 knockdown reduced levels of immunoproteasome-specific subunits, such as PSMB8 (β-5i) and PA28a compared with control cells. Of note, PI31 knockdown did not alter the expression of constitutive proteasome subunits, such as PSMD2, PSMA4 (α4) and PMSA2 (α2) (fig. 3I). Together these results show a previously unidentified cell intrinsic role for PI31 in immune cells to modulate the generation of HLA class I-associated peptides, partially through stabilising immunoproteasome subunit composition.

### SEC16A forms a complex with PI31 and TBK1

To identify interaction partners of PI31, we immunoprecipitated PI31 from NOD2 and TLR2 stimulated DCs, followed by performing tryptic digestion and mass spectrometry analysis (fig. 4A). After NOD2 and TLR2 stimulation, we detected SEC16A as the protein with the highest confidence score and number of unique peptides (fig. 4B, C). SEC16A is an endoplasmic reticulum (ER) protein that marks ER exit sites (ERES) in mammalian cells and is implicated in ER to Golgi vesicular trafficking (42), so we sought to confirm an interaction between PI31 and SEC16A. We immunoprecipitated PI31 in both primary DCs and THP1 cells following NOD2 activation and examined SEC16A levels by western blot. This result confirmed the presence of PI31 and SEC16A in the same protein complex in both non-stimulated and stimulated samples, though increased SEC16A levels occurred in samples stimulated with MDP and PAM_3_CSK_4_ (fig. 4D, E). We then overexpressed GFP-tagged PI31 in THP1 cells and assessed co-localization with SEC16A by confocal microscopy. We observed a significantly enriched co-localization of PI31 and SEC16A, which was dependent on NOD2 and TLR2 signals in DCs (fig. 4F, G). Given the direct association of PI31 with TBK1, we tested whether SEC16A and TBK1 are present in the same protein complex. So, we immunoprecipitated endogenous TBK1 in THP1 cells and examined levels of SEC16A by western blot. Co-immunoprecipitation confirmed the association of TBK1 and SEC16A in the same protein complex (fig. 4H).

**Figure 4.**
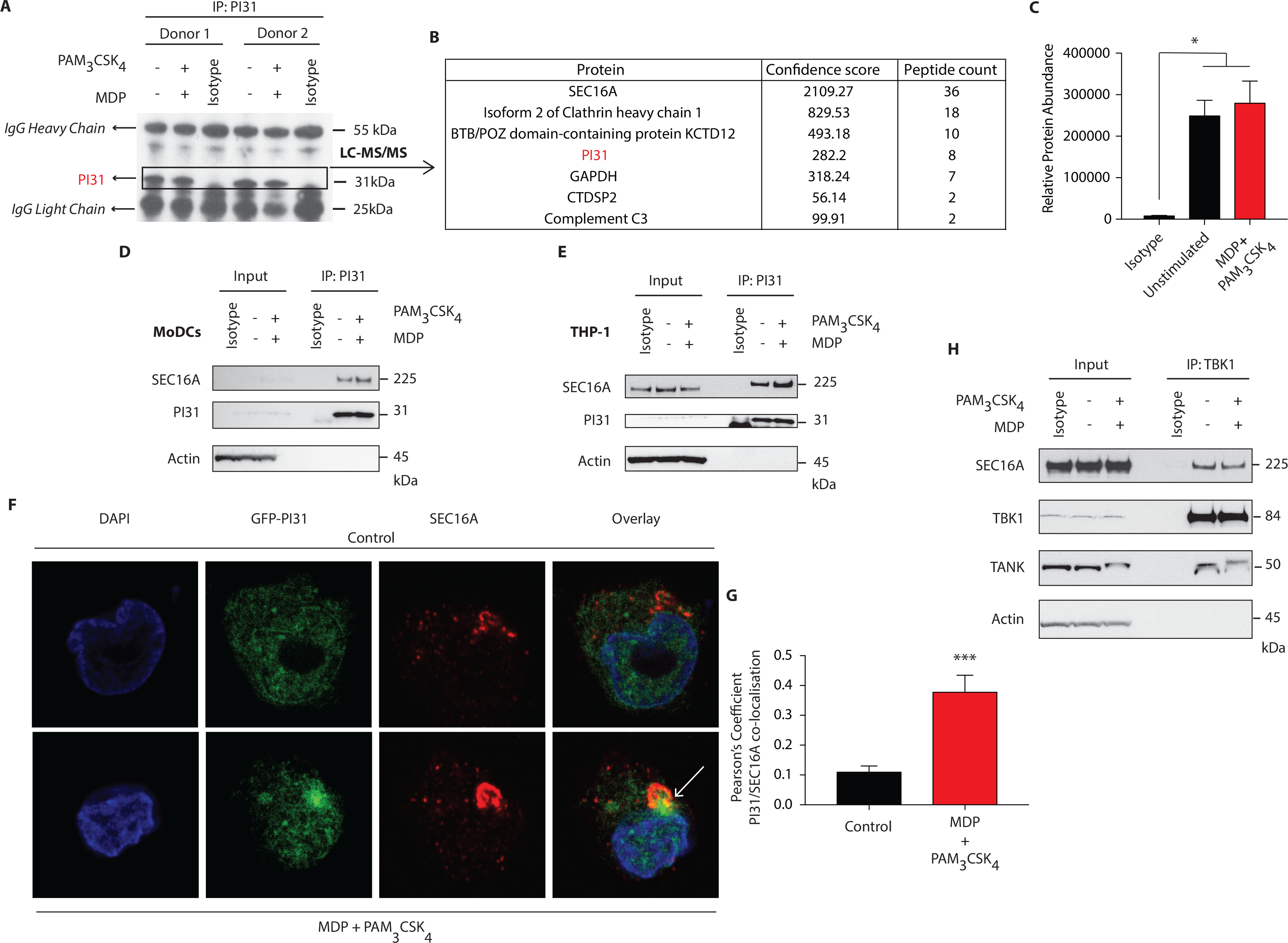
SEC16A forms a complex with PI31 and TBK1. (**A**) Immunoblot analysis representing immunoprecipitation of endogenous PI31 in DCs following MDP/PAM_3_CSK_4_ stimulation for LC-MS/MS analysis. (**B**) List of proteins from LC/MS analysis with highest confidence score and peptide counts. (**C**) Relative abundance protein for SEC16A. Data represent mean ± s.e.m; (n=2) \**P*<0.05 Student’s *t* test. (**D**, **E**) DCs (**D**) and THP1 (**E**) cells stimulated with MDP/PAM_3_CSK_4_. Cell lysates were immunoprecipitated with PI31 antibodies and probed with anti-SEC16A. Data are representative of 3 independent experiments. (**F**, **G**) Images show confocal analysis of THP-1 cells transfected with GFP-PI31 (green) for 24 hours, and stimulated with MDP/PAM_3_CSK_4_ for 30 minutes and stained with antibody to SEC16A (red) and DAPI (blue). Pearson’s coefficient for PI31 and SEC16A co-localization in control or MDP/PAM_3_CSK_4_ stimulated cells. Data represent means ± s.e.m. (n=20), Three independent experiments with at least five images analyzed per experiment); \*\*\**P*<0.0001 Student’s *t* test. (**H**) THP1 cells were stimulated with MDP/PAM_3_CSK_4_ for 30 min. Cell lysates were immunoprecipitated with anti-TBK1 antibodies and probed with anti-SEC16A, anti-TBK1 and anti-TANK antibodies. Data represent 3 independent experiments.

As SEC16A is a protein implicated in vesicular trafficking from the ER, we next examined whether NOD2 and TLR2 stimulation could control SEC16A recruitment to phagosomes carrying components of PGN. We treated BMDCs from WT mice with streptavidin beads conjugated to biotinylated MDP and PAM_3_CSK_4_ and examined cellular localization of SEC16A by confocal microscopy. Phagosomes bearing beads with or without MDP and PAM_3_CSK_4_ failed to acquire SEC16A (fig. S4A). We confirmed these findings by examining LAMP1 levels by immunoblot following stimulation of BMDCs with streptavidin beads conjugated to biotinylated MDP, PAM_3_CSK_4_ and LPS. While SEC16A, TBK1 and PI31 were not enriched in phagosomes carrying beads, they were enriched only in whole cell extracts (WCE) when the ER resident ERp72 was detected. Phagosomal lysates showed enriched LAMP-1 with significantly increased levels on phagosome carrying beads conjugated to MDP/PAM_3_CSK_4_ and LPS (fig. S4B).

These results indicate a novel association between PI31 and SEC16A, a component of endoplasmic reticulum exit sites (ERES), required for secretory cargo traffic from the ER to the Golgi apparatus. We demonstrated that the levels of PI31 and SEC16A in this complex are increased on NOD2 and TLR2 sensing. We further demonstrated the association of TBK1 in the same protein complex.

### PI31 modulate NOD2/TLR2-dependent cross-presentation and CD8^+^ T cell activation

Studies demonstrated that PRR signaling, including NOD2, increases CD8^+^ T cell activation by enhancing cross-presentation (43,27,44). However, the mechanisms underlying the effects of PRR engagement on antigen presentation are incompletely understood. We determined the impact of NOD2 and TLR2 dependent phosphorylation of PI31 on cross-presentation by downregulating *PI31* expression by siRNA knockdown in DCs derived from the bone marrow (BMDC) from wild-type (WT) mice stimulated with MDP and PAM_3_CSK_4_ and pulsed with full length ovalbumin (OVA) (fig. 5A) and quantified CD8^+^ T cell activation through increased CD69 expression. We found significantly reduced cross-presentation of OVA to OT-I T cells in BMDCs expressing low levels of PI31 compared to cells expressing PI31 (fig. 5B). In contrast, we found the octapeptide SIINFEKL (OVA amino acids 257–264), which is directly recognized by OT-I T cells and does not require proteasomal degradation, was presented with equal efficiency by BMDCs expressing normal or reduced levels of PI31 (fig. 5C). These data indicate that cross-presentation requires PI31 in DCs stimulated with NOD2 and TLR2 ligands.

**Figure 5.**
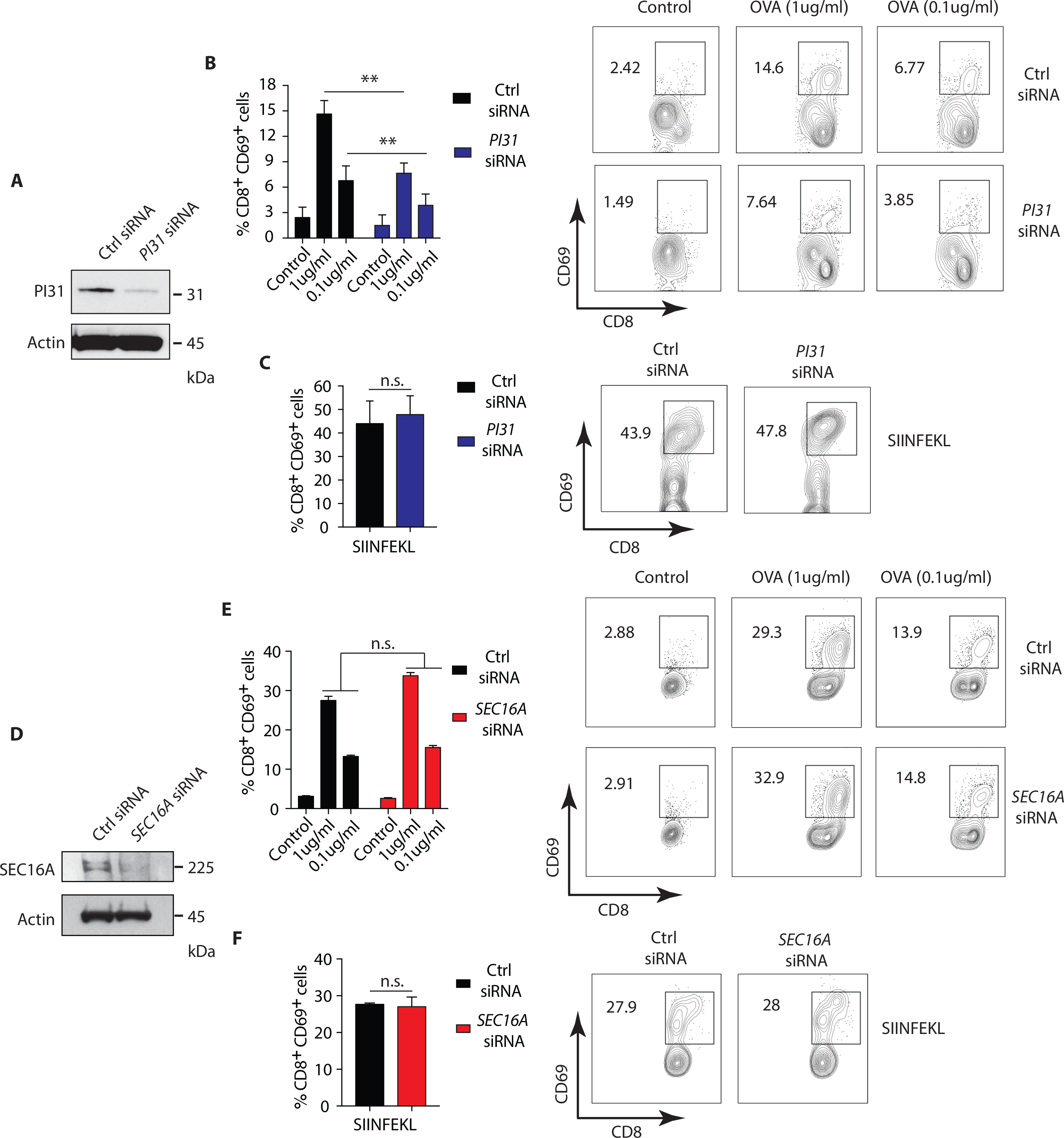
PI31, but not SEC16A, mediates NOD2/TLR2-dependent cross-presentation. (**A**) PI31 expression was downregulated by transfecting BMDCs with control non-targeting and *PI31* siRNAs. Immunoblots data represent 3 independent experiments. (**B**, **C)** 48 hours post-transfection, BMDCs were incubated with MDP/PAM_3_CSK_4_ in presence or absence of sOVA (1 and 0.1ug/ml) or SIINFEKL peptide for 3 hours followed by co-culture with OT-I CD8^+^ T cells for 72 hrs. Cross-presentation efficiency of DCs was determined by the percentage of CD8^+^CD69^+^ T cells using FACS. Data represent the mean ± s.e.m; (B, C, n=4) \*\**P*<0.01 Student’s *t* test. Data are from one experiment representative of three or more independent experiments. (**D**) SEC16A expression was downregulated by transfecting BMDCs with control non-targeting and *SEC16A* siRNAs. Immunoblots data represent 3 independent experiments. (**E**, **F**) 48 hours post-transfection, BMDCs were incubated with MDP/PAM_3_CSK_4_ with or without sOVA (1 and 0.1ug/ml) or SIINFEKL peptide for 3 hours followed by co-culture with OT-I CD8^+^ T cells for 72 hrs. Cross-presentation was measured by calculating the percentage of CD8^+^CD69^+^ T cells using FACS. Data represent mean ± s.e.m; (E, F, n=3). Data from one experiment representative of three or more independent experiments.

Then, we investigated whether the PI31 binding partner, SEC16A, also increased cross-presentation during stimulation with NOD2 and TLR2. We knocked down *SEC16A* expression in BMDCs (fig. 5D), followed by stimulation with MDP and PAM_3_CSK_4_ plus ovalbumin (OVA). We assessed cross-presentation to CD8^+^ TCR transgenic OT-I T cells as previously described. We found no change in cross-presentation of OVA to OT-I T cells in BMDCs expressing low levels of SEC16A compared to cells expressing SEC16A (fig. 5E). We found an equal efficiency in presentation of SIIFENKL by BMDCs expressing normal or reduced PI31 levels (fig. 5F). Although SEC16A forms a complex with PI31 on NOD2 triggering, these results suggest that it is not required to modulate cross-presentation. Thus, NOD2/TLR2 mediated MHC class I antigen presentation requires PI31, but not SEC16A, *in vitro*.

### TBK1 inhibition prevents cross-presentation

To investigate whether the involvement of NOD2/TLR2-dependent cross-presentation requires TBK1 activity, we pretreated BMDCs from WT mice with the TBK1 specific inhibitor, BX795, followed by stimulation with MDP and PAM_3_CSK_4_ and OVA. Treatment with BX795 significantly reduced cross-presentation of OVA to OT-I T cells in BMDCs compared to control cells (fig. 6A), while the octapeptide SIINFEKL was presented with equal efficiency by MDP and PAM_3_CSK_4_ stimulated BMDCs with or without BX795 (fig. 6B). Then, we investigated whether NOD2/TLR2-dependent TBK1 activity also promoted cross-presentation in DCs *in vivo*. We treated WT mice with Amlexanox, which blocks TBK1 activity *in vivo* without affecting the activity of IKK-α or IKK-β or any other known kinases (45,46). We administered Amlexanox intraperitoneally (i.p.) and administered a second injection of the inhibitor after 24 hrs accompanied by i.p. injection of MDP and PAM_3_CSK_4_. After 12 hrs, we injected mice with OVA, isolated total splenic CD11c^+^ cells and co-cultured them with OT-I CD8^+^ T cells to measure antigen-specific T cell activation (fig. 6C).

**Figure 6.**
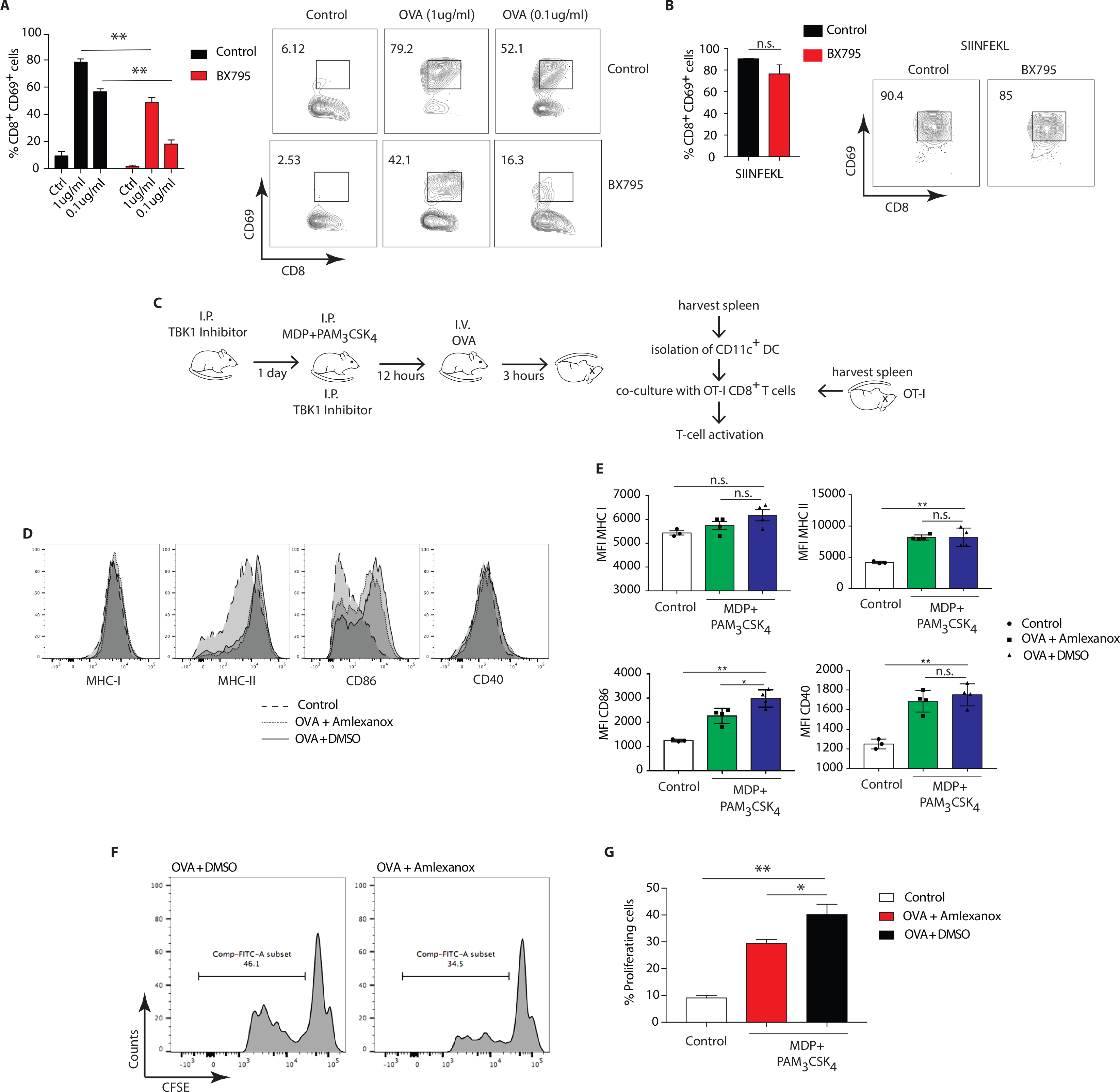
TBK1 mediates NOD2/TLR2-dependent cross-presentation. (**A**, **B**) BMDCs were treated for 1 hr with BX795 (1 μM). BMDCs were then incubated with MDP/PAM_3_CSK_4_ with or without sOVA (1 and 0.1ug/ml) or SIINFEKL peptide for 3 hrs prior to co-culture with OT-I CD8^+^ T cells for 72 hrs. Cross-presentation was measured by calculating the percentage of CD8^+^CD69^+^ T cells using FACS. Data represent the mean ± s.e.m (n=3) \*\**P*<0.01 Student’s *t* test. Data represent one experiment representative of three independent experiments. (**C**) Experimental scheme for *in vivo* cross-presentation assay. (**D**) FACS analysis indicating levels of MHC-I, MHC-II, CD86 and CD40 on DCs treated as in C. (**E**) MFI values relative to MHC-I, MHC-II, CD86 and CD40 show DC activation levels. Data indicate mean ± s.e.m. (*n*=3; *n*=4). n.s. = non-significant; ** *P*<0.01; * *P*<0.05; one-way ANOVA (Tukey’s multiple comparison test). (**F**) Efficiency of cross-presentation measured by proliferation of CFSE-labeled OT-I CD8^+^ T cells and FACS analysis of CFSE fluorescence, indicating level of OT-I CD8^+^ T cell activation. (**G**) T cell proliferation status determined by FACS and plotted as the percentage of dividing OT-I-ovalbumin-reactive CD8^+^ T cells. Data represent mean ± s.e.m. (*n*=8); ** *P*<0.01; * *P*<0.05; one-way ANOVA (Tukey’s multiple comparison test). Data are from one experiment representative of two independent experiments.

First, we tested whether stimulation with MDP and PAM_3_CSK_4_ enhanced activation of DCs by assessing expression of DC activation markers. NOD2 and TLR2 stimulation significantly increased expression of MHC-II, CD86, CD40, but not MHC-I, compared to control mice. Treatment with Amlexanox did not change the expression of MHC-II, CD40 and MHC-I (fig. 6D, E). We assessed OT-I CD8^+^ T cell specific activation by examining the percentage of proliferation measured by CFSE dye dilution. Of note, DCs from mice pre-treated with Amlexanox cross-presented OVA significantly less than untreated mice (fig. 6F, G). We conclude that enhanced MHC class I cross presentation mediated by DCs activated *in vivo* with MDP and PAM_3_CSK_4_ requires TBK1 activity.

### DCs expressing *NOD2* variants from Crohn’s disease show defective cross-presentation

Polymorphisms in *NOD2* represent the strongest known genetic risk factors associated with the development of Crohn’s disease (47). So, we investigated whether Crohn’s patients DCs expressing variants of NOD2 exhibited defects in cross-presentation after NOD2 stimulation. We used HLA-A2 DCs from healthy individuals expressing wild type (WT) *NOD2* or HLA-A2 Crohn’s patients DCs expressing Crohn’s associated *NOD2* polymorphisms. We assessed the cross-presentation ability of DCs using the NY-ESO-1 specific CD8^+^ CTL clones generated by directly sorting HLA-A2 NY-ESO-1 _157-165_ tetramers from a melanoma patient (48,28).

Following the expansion of the CTL clones, we stimulated HLA-A2^+^ patients DCs either homozygous for 1007fsinsC NOD2 expression or compound heterozygous for any Crohn’s-associated NOD2 polymorphisms with MDP or MDP+ PAM_3_CSK_4_ in the presence of the 19-mer NY-ESO-1_151-169_ peptide (SCLQQLSLLMWITQCFLPV) or with the shorter NY-ESO-1_157-165_ 9- mer peptide (SLLMWITQC) directly binding to HLA-A2 MHC class I molecules (fig. 7A). We determined cross-presentation following co-culture of DCs and NY-ESO-1 specific CD8^+^ CTL clones for 24 hrs and assessed the percentage of CD44^+^CD25^+^ CD8^+^ T cells present. We found decreased cross-presentation of the NY-ESO-1_151-169_ peptide (19-mer) to CD8^+^ T cells in Crohn’s DCs expressing *NOD2* polymorphisms compared to DCs from healthy donors with WT *NOD2* (fig. 7B, C) following stimualtion with MDP or MDP+PAM_3_CSK_4_. In contrast, we found no significant difference in T cell responses to the NY-ESO-1_157-165_ peptide (9-mer) that directly bound to surface HLA-A2 MHC class I molecules between healthy or Crohn’s DCs stimulated with MDP (fig. 7D, E). Thus, our data indicate that Crohn’s donor DCs show defective cross-presentation following NOD2 stimulation and may indicate aberrant intestinal CD8^+^ T cell responses detected in the mucosa in Crohn’s disease.

**Figure 7.**
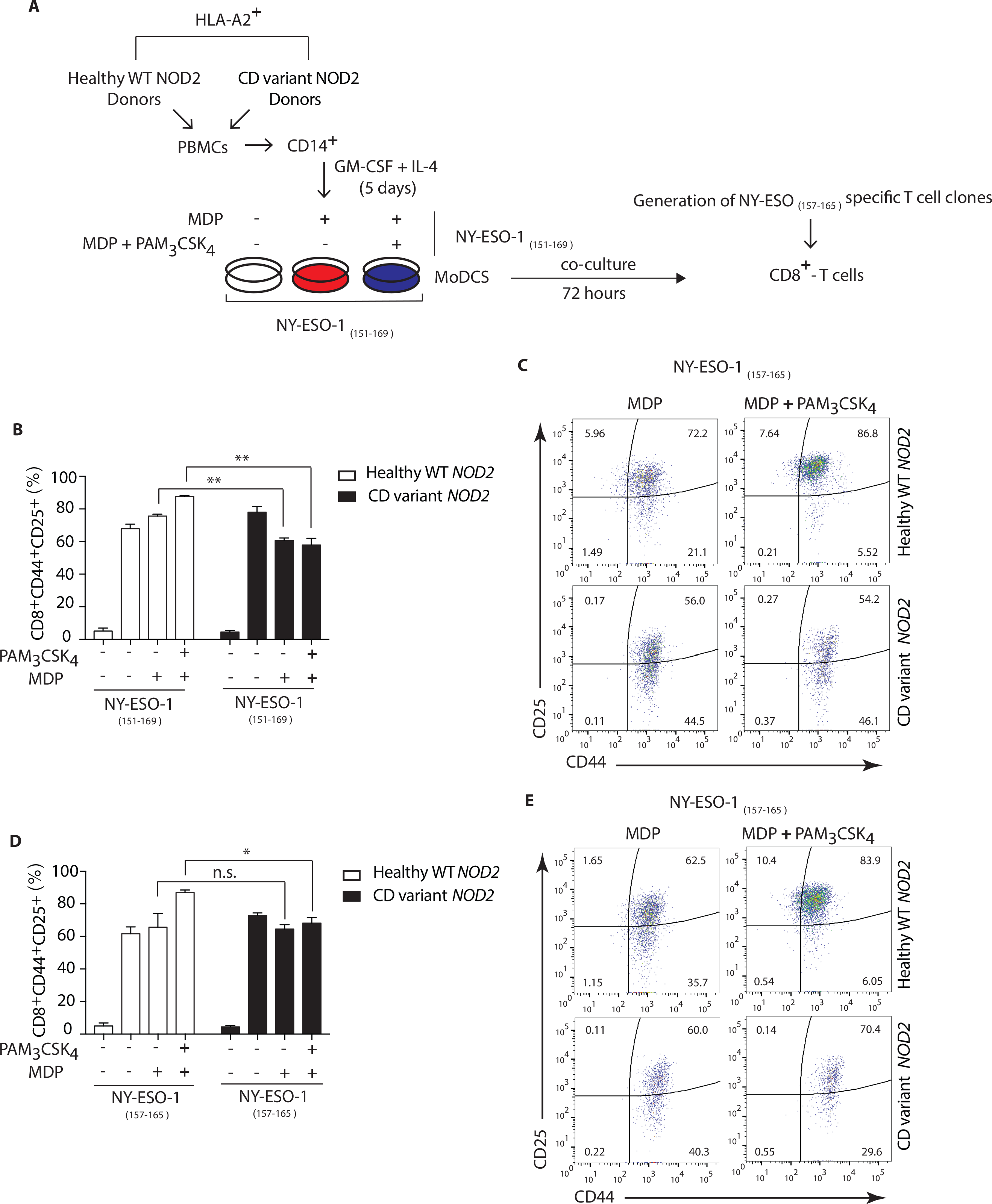
Crohn’s patients DCs show impaired NOD2/TLR2-dependent cross-presentation. (**A**) DCs were derived from HLA-A2^+^ Crohn’s patients homozygous for FS1007insC NOD2 or R702W + G908R NOD2 compound heterozygotes or healthy HLA-A2^+^ volunteers expressing wild type NOD2. HLA-A2^+^ DCs from healthy donors or Crohn’s patients were incubated with MDP or MDP + PAM_3_CSK_4_ in the presence of either the 19-mer NY-ESO-1_151-169_ peptide (SCLQQLSLLMWITQCFLPV) or with the shorter 9-mer NY-ESO-1_157-165_ peptide (SLLMWITQC) for 3 hrs followed by co-culture with the NY-ESO-1_157-165_ specific CD8^+^ CTL clones for 24 hrs. (**B, C)** MHC class I cross-presentation of 19-mer NY-ESO-1_151-169_ peptides by DCs was measured by calculating the percentage of CD8^+^CD44^+^CD25^+^ T cells using FACS analysis. (**D, E**) MHC class I direct presentation of 9-mer NY-ESO-1_157-165_ peptides by DCs was measured by calculating the percentage of CD8^+^CD44^+^CD25^+^ T cells using FACS analysis. Data represent mean ± s.e.m. (*n*=6); ** *P*<0.01; * *P*<0.05; one-way ANOVA (Tukey’s multiple comparison test).

## Discussion

Here, we reveal that NOD2 and TLR2 signal to PI31 via TBK1 to facilitate MHC-I antigen presentation in DCs and prime CD8^+^ T cell responses. TANK-binding kinase-1 (TBK1) mediates this effect by modulating PI31 phosphorylation following MDP and PAM_3_CSK_4_ stimulation and directly associating with PI31. By undertaking a mass spectrometry analysis of MHC-I/peptide complexes expressed on DCs, we identified an unknown intrinsic function of PI31 in regulating the quantity of peptides associated with MHC-I, likely through stabilizing the immunoproteasome. We discovered an interaction between PI31 and SEC16A, a component of endoplasmic reticulum exit sites (ERES), which forms a complex with TBK1 and PI31 increased by MDP and PAM_3_CSK_4_ exposure. Depleting PI31 or inhibiting TBK1 activity in DCs stimulated with NOD2 and TLR2 ligands impaired cross-presentation and CD8^+^ T cell activation. Finally, we examined DCs from patients with Crohn’s disease expressing associated *NOD2* variants and found they were unable to induce cross-presentation following NOD2 and TLR2 triggering.

PI31 remains the most poorly characterized proteasome regulator. The C-terminal domain of PI31, which possesses an intrinsically disordered structure, was originally characterized as an *in vitro* inhibitor of 20S proteasome activity (49-52). However, this C-terminal site contains an HbY*X* motif characteristic of multiple proteasome activators. Consistent with this finding, this PI31 activates the 26S proteasome *in vitro* and exerts a positive effect on proteasome function in intact cells (52). These contradictory findings are the focus of investigations by Li et al., who found multiple regions in PI31 bind independently to the proteasome to exert specific actions *in vitro* (53). However, they detected no change in cellular constitutive proteasome function with increased or decreased PI31 levels by overexpression or RNA interference (RNAi) respectively (53). So, despite *in vitro* effects on various proteasome activities, the cellular roles and mechanisms of PI31 in regulating proteasome function remain unclear. Our results presented here further elucidate PI31 function demonstrating an unidentified role in antigen presenting cells directly modulating the generation of HLA class I-associated peptides partially through stabilising immunoproteasome subunit composition.

The interplay between NOD2 and TLR2 has been well characterized and it is not surprising given that both receptors respond to adjacent components of PGN expressed on bacterial cell walls, MDP and PAM_3_CSK_4_ respectively (10,6,7,11,54,9). Upon MDP recognition, NOD2 recruits RIPK2, which induces activation of the IKK complex. IKK phosphorylates IκBα, targeting it for Lys48-linked polyubiquitination and proteasome-dependent degradation facilitating NF-κB translocation into the nucleus (55,31). This signaling pathway, in physiological conditions, amplifies TLR2 signals and both synergize in the induction of NF-κB, which play a central role in cytokine production (56,7,57,58). PI31 did not affect either NF-κB activation or proteasome-dependent degradation of IκBα mediated by NOD2/TLR2. NOD2 induces formation of a protein complex containing RIPK2 and TRAF3, which recruits TBK1 to induce type I IFN (36). TBK-1 is a member of the IKK family of kinases falling into the category of the non-canonical IKK-related kinases (IKKε and TBK-1) (59). We found TBK1 directly associates with PI31 to direct phosphorylation on Serine 153 following NOD2 and TLR2 triggering and this represents a novel signaling pathway directing subsequent MHC I antigen presentation on PGN recognition. PI31 and TBK1 had no effect on the direct presentation of the octapeptide SIINFEKL (OVA amino acids 257–264). OT-I T cells directly recognize SIINFEKL and do not require proteasomal degradation excluding a role for TBK1 mediated type I IFN production in inducing CD8^+^ T cell responses.

Several studies demonstrated that PRR engagement and simultaneous exposure to foreign antigen increases CD8^+^ T cell activation by cross-presented peptides (25). Different proposed mechanisms explain the increased cross-presentation observed during the initial phase of TLR sensing in immature DCs. These include enhancing recruitment of MHC class I to phagosomes from endosomal recycling compartments (ERC) (27), maintaining low levels of lysosomal proteases (60) or recruiting the NADPH oxidase NOX2 to early phagosomes for sustained production of low levels of reactive oxygen species and phagosomal alkalinization (61). Whether distinctions occur through which the mechanisms dictating which different classes of PRR interact with the proteasome is unclear. NOD2 can modulate cross-presentation, and our results are consistent with a study demonstrating that NOD2 signals significantly augment the cross-priming of Ag-specific CD8^+^ T cells *in vivo* by upregulating MHC class I-dependent Ag presentation pathway (43). During late phases of DC maturation, cross-presentation is down-modulated (62-64). For example, after 24–30 hrs of LPS stimulation, DCs do not cross-present antigens, likely through reducing antigen export to the cytosol (44). In addition, pre-treatment with pure PGN, as well as both synthetic ligands for NOD1 and NOD2, impaired cross-presentation of HSV Ags or of vaccinia virus-expressed OVA (VV-OVA) after 22 hr pre-treatment (63). Thus, early non-sustained signaling events, such as TBK1 mediated PI31 phosphorylation driven by PRRs, may indicate temporal control of MHC I antigen presentation and prevent excessive priming during chronic phases of pathogen handling. We found PI31 forms a complex with both SEC16A and TBK1 on NOD2/TLR2 stimulation that localized at the ER but not phagosomal contact sites. SEC16A marks ER exit sites (ERES) and directs ER to Golgi transport(65) and may contribute to recruitment of the peptide loading complex to ER-Golgi intermediated compartment (ERGIC). While SEC16A could enable egress of the peptide loading complex, functional redundancy most likely occurs, as we found no effect of reduced SEC16A levels by siRNA on antigen cross-presentation *in vitro*.

*NOD2* remains the most strongly associated Crohn’s disease susceptibility gene (66,67). *NOD2* polymorphisms (R702W, G908R, and 1007fs) in Crohn’s are often associated with disease states affecting the terminal ileum and contributing to stricturing and fibrostenosing disease requiring surgical intervention (68). While defective autophagy occurring in the presence of Crohn’s associated NOD2 leads to aberrant MHC class II responses (13), it is unclear whether Crohn’s associated *NOD2* polymorphisms also affect MHC class I responses. Our work elucidates an unidentified immune function for NOD2 in human DCs. Our results demonstrate that NOD2 via the cross-presentation pathway drives CD8^+^ T cell activation and in the presence of Crohn’s associated *NOD2* polymorphisms impairs these responses. As a result of impaired T cell mediated clearance of commensal microbes, we expect loss of function NOD2 mutations to impair antigen presentation leading to non-specific intestinal inflammation. The adjuvant activity of MDP and TLR ligands is well known to enhance vaccine responses (69). Here, we define the molecular basis for this adjuvancy. We speculate that biochemically disparate forms of PGN derived from different bacterial species may affect immunoproteasome stability and antigen presentation to different degrees.

## Acknowledgments

We acknowledge support of the Oxford NIHR Biomedical Research Centre. Mass spectrometry sample acquisition was performed in the Target Discovery Institute, Mass Spectrometry Laboratory led by Professor Benedikt Kessler. The authors acknowledge the Wolfson Imaging Centre, Weatherall Institute of Molecular Medicine (Oxford, UK), Helen Ferry at the Experimental Medicine Division Flow Cytometry Facility, and the NIHR IBD Bioresource for Identification of Crohn’s patients expressing specific NOD2 polymorphisms.

## Author contributions

D.C. and A.S. designed experimental studies, interpreted the data and wrote the manuscript. D.C., S.S., T.C., T.S., D.M., M.L.T., G.P., JL.C. U.G. and N.T. performed experiments, acquired and analyzed data. V.C. provided critical reagents, helped interpret data and edit the manuscript. A. S. supervised and obtained funding.

## References

Akira, S, Uematsu, S, and Takeuchi, O (2006). Pathogen recognition and innate immunity. Cell 124, 783–801. doi: 10.1016/j.cell.2006.02.015.

Caruso, R, Warner, N, Inohara, N, and Nunez, G (2014). NOD1 and NOD2: signaling, host defense, and inflammatory disease. Immunity 41, 898–908. doi: 10.1016/j.immuni.2014.12.010.

Ferwerda, G, Girardin, SE, Kullberg, BJ, Le Bourhis, L, de Jong, DJ, Langenberg, DM, et al. (2005). NOD2 and toll-like receptors are nonredundant recognition systems of Mycobacterium tuberculosis. PLoS Pathog 1, 279–285. doi: 10.1371/journal.ppat.0010034.

Hemmi, H, and Akira, S (2005). TLR signalling and the function of dendritic cells. Chem Immunol Allergy 86, 120–135. doi: 10.1159/000086657.

Kufer, TA, Kremmer, E, Banks, DJ, and Philpott, DJ (2006). Role for erbin in bacterial activation of Nod2. Infect Immun 74, 3115–3124. doi: 10.1128/IAI.00035-06.

Kobayashi, KS, Chamaillard, M, Ogura, Y, Henegariu, O, Inohara, N, Nunez, G, et al. (2005). Nod2-dependent regulation of innate and adaptive immunity in the intestinal tract. Science 307, 731–734. doi: 10.1126/science.1104911.

Netea, MG, Ferwerda, G, de Jong, DJ, Jansen, T, Jacobs, L, Kramer, M, et al. (2005). Nucleotide-binding oligomerization domain-2 modulates specific TLR pathways for the induction of cytokine release. J Immunol 174, 6518–6523.

Philpott, DJ, and Girardin, SE (2004). The role of Toll-like receptors and Nod proteins in bacterial infection. Mol Immunol 41, 1099–1108. doi: 10.1016/j.molimm.2004.06.012.

Kim, YG, Park, JH, Shaw, MH, Franchi, L, Inohara, N, and Nunez, G (2008). The cytosolic sensors Nod1 and Nod2 are critical for bacterial recognition and host defense after exposure to Toll-like receptor ligands. Immunity 28, 246–257. doi: 10.1016/j.immuni.2007.12.012.

Netea, MG, Kullberg, BJ, de Jong, DJ, Franke, B, Sprong, T, Naber, TH, et al. (2004). NOD2 mediates anti-inflammatory signals induced by TLR2 ligands: implications for Crohn’s disease. Eur J Immunol 34, 2052–2059. doi: 10.1002/eji.200425229.

van Heel, DA, Ghosh, S, Butler, M, Hunt, KA, Lundberg, AM, Ahmad, T, et al. (2005). Muramyl dipeptide and toll-like receptor sensitivity in NOD2-associated Crohn’s disease. Lancet 365, 1794–1796. doi: 10.1016/S0140-6736(05)66582-8.

Delgado, MA, Elmaoued, RA, Davis, AS, Kyei, G, and Deretic, V (2008). Toll-like receptors control autophagy. EMBO J 27, 1110–1121. doi: 10.1038/emboj.2008.31.

Cooney, R, Baker, J, Brain, O, Danis, B, Pichulik, T, Allan, P, et al. (2010). NOD2 stimulation induces autophagy in dendritic cells influencing bacterial handling and antigen presentation. Nat Med 16, 90–97. doi: 10.1038/nm.2069.

Travassos, LH, Carneiro, LA, Ramjeet, M, Hussey, S, Kim, YG, Magalhaes, JG, et al. (2010). Nod1 and Nod2 direct autophagy by recruiting ATG16L1 to the plasma membrane at the site of bacterial entry. Nat Immunol 11, 55–62. doi: 10.1038/ni.1823.

van Beelen, AJ, Zelinkova, Z, Taanman-Kueter, EW, Muller, FJ, Hommes, DW, Zaat, SA, et al. (2007). Stimulation of the intracellular bacterial sensor NOD2 programs dendritic cells to promote interleukin-17 production in human memory T cells. Immunity 27, 660–669. doi: 10.1016/j.immuni.2007.08.013.

Magalhaes, JG, Rubino, SJ, Travassos, LH, Le Bourhis, L, Duan, W, Sellge, G, et al. (2011). Nucleotide oligomerization domain-containing proteins instruct T cell helper type 2 immunity through stromal activation. Proc Natl Acad Sci U S A 108, 14896–14901. doi: 10.1073/pnas.1015063108.

Brain, O, Owens, BM, Pichulik, T, Allan, P, Khatamzas, E, Leslie, A, et al. (2013). The intracellular sensor NOD2 induces microRNA-29 expression in human dendritic cells to limit IL-23 release. Immunity 39, 521–536. doi: 10.1016/j.immuni.2013.08.035.

Corridoni, D, Rodriguez-Palacios, A, Di Stefano, G, Di Martino, L, Antonopoulos, DA, Chang, EB, et al. (2017). Genetic deletion of the bacterial sensor NOD2 improves murine Crohn’s disease-like ileitis independent of functional dysbiosis. Mucosal Immunol 10, 971–982. doi: 10.1038/mi.2016.98.

Joffre, OP, Segura, E, Savina, A, and Amigorena, S (2012). Cross-presentation by dendritic cells. Nat Rev Immunol 12, 557–569. doi: 10.1038/nri3254.

Palmowski, MJ, Gileadi, U, Salio, M, Gallimore, A, Millrain, M, James, E, et al. (2006). Role of immunoproteasomes in cross-presentation. J Immunol 177, 983–990.

Shen, L, Sigal, LJ, Boes, M, and Rock, KL (2004). Important role of cathepsin S in generating peptides for TAP-independent MHC class I crosspresentation in vivo. Immunity 21, 155–165. doi: 10.1016/j.immuni.2004.07.004.

Beutler, B, Hoebe, K, Du, X, and Ulevitch, RJ (2003). How we detect microbes and respond to them: the Toll-like receptors and their transducers. J Leukoc Biol 74, 479–485. doi: 10.1189/jlb.0203082.

Kopp, E, and Medzhitov, R (2003). Recognition of microbial infection by Toll-like receptors. Curr Opin Immunol 15, 396–401.

Burgdorf, S, Scholz, C, Kautz, A, Tampe, R, and Kurts, C (2008). Spatial and mechanistic separation of cross-presentation and endogenous antigen presentation. Nat Immunol 9, 558–566. doi: 10.1038/ni.1601.

Blander, JM (2016). The comings and goings of MHC class I molecules herald a new dawn in cross-presentation. Immunol Rev 272, 65–79. doi: 10.1111/imr.12428.

Corridoni, D, and Simmons, A (2017). Innate immune receptors for cross-presentation: The expanding role of NLRs. Mol Immunol. doi: 10.1016/j.molimm.2017.11.028.

Nair-Gupta, P, Baccarini, A, Tung, N, Seyffer, F, Florey, O, Huang, Y, et al. (2014). TLR signals induce phagosomal MHC-I delivery from the endosomal recycling compartment to allow cross-presentation. Cell 158, 506–521. doi: 10.1016/j.cell.2014.04.054.

Chen, JL, Morgan, AJ, Stewart-Jones, G, Shepherd, D, Bossi, G, Wooldridge, L, et al. (2010). Ca2+ release from the endoplasmic reticulum of NY-ESO-1-specific T cells is modulated by the affinity of TCR and by the use of the CD8 coreceptor. J Immunol 184, 1829–1839. doi: 10.4049/jimmunol.0902103.

Zaiss, DM, Standera, S, Kloetzel, PM, and Sijts, AJ (2002). PI31 is a modulator of proteasome formation and antigen processing. Proc Natl Acad Sci U S A 99, 14344–14349. doi: 10.1073/pnas.212257299.

Cho-Park, PF, and Steller, H (2013). Proteasome regulation by ADP-ribosylation. Cell 153, 614–627. doi: 10.1016/j.cell.2013.03.040.

Hasegawa, M, Fujimoto, Y, Lucas, PC, Nakano, H, Fukase, K, Nunez, G, et al. (2008). A critical role of RICK/RIP2 polyubiquitination in Nod-induced NF-kappaB activation. EMBO J 27, 373–383. doi: 10.1038/sj.emboj.7601962.

Canning, P, Ruan, Q, Schwerd, T, Hrdinka, M, Maki, JL, Saleh, D, et al. (2015). Inflammatory Signaling by NOD-RIPK2 Is Inhibited by Clinically Relevant Type II Kinase Inhibitors. Chem Biol 22, 1174–1184. doi: 10.1016/j.chembiol.2015.07.017.

Nachbur, U, Stafford, CA, Bankovacki, A, Zhan, Y, Lindqvist, LM, Fiil, BK, et al. (2015). A RIPK2 inhibitor delays NOD signalling events yet prevents inflammatory cytokine production. Nat Commun 6, 6442. doi: 10.1038/ncomms7442.

Ogura, Y, Inohara, N, Benito, A, Chen, FF, Yamaoka, S, and Nunez, G (2001). Nod2, a Nod1/Apaf-1 family member that is restricted to monocytes and activates NF-kappaB. J Biol Chem 276, 4812–4818. doi: 10.1074/jbc.M008072200.

Watanabe, T, Asano, N, Fichtner-Feigl, S, Gorelick, PL, Tsuji, Y, Matsumoto, Y, et al. (2010). NOD1 contributes to mouse host defense against Helicobacter pylori via induction of type I IFN and activation of the ISGF3 signaling pathway. J Clin Invest 120, 1645–1662. doi: 10.1172/JCI39481.

Correa, RG, Milutinovic, S, and Reed, JC (2012). Roles of NOD1 (NLRC1) and NOD2 (NLRC2) in innate immunity and inflammatory diseases. Biosci Rep 32, 597–608. doi: 10.1042/BSR20120055.

Clark, K, Plater, L, Peggie, M, and Cohen, P (2009). Use of the pharmacological inhibitor BX795 to study the regulation and physiological roles of TBK1 and IkappaB kinase epsilon: a distinct upstream kinase mediates Ser-172 phosphorylation and activation. J Biol Chem 284, 14136–14146. doi: 10.1074/jbc.M109.000414.

Cerundolo, V, Kelly, A, Elliott, T, Trowsdale, J, and Townsend, A (1995). Genes encoded in the major histocompatibility complex affecting the generation of peptides for TAP transport. Eur J Immunol 25, 554–562. doi: 10.1002/eji.1830250238.

Cerundolo, V, Benham, A, Braud, V, Mukherjee, S, Gould, K, Macino, B, et al. (1997). The proteasome-specific inhibitor lactacystin blocks presentation of cytotoxic T lymphocyte epitopes in human and murine cells. Eur J Immunol 27, 336–341. doi: 10.1002/eji.1830270148.

Neefjes, J, Jongsma, ML, Paul, P, and Bakke, O (2011). Towards a systems understanding of MHC class I and MHC class II antigen presentation. Nat Rev Immunol 11, 823–836. doi: 10.1038/nri3084.

Rock, KL, Gramm, C, Rothstein, L, Clark, K, Stein, R, Dick, L, et al. (1994). Inhibitors of the proteasome block the degradation of most cell proteins and the generation of peptides presented on MHC class I molecules. Cell 78, 761–771.

Hughes, H, Budnik, A, Schmidt, K, Palmer, KJ, Mantell, J, Noakes, C, et al. (2009). Organisation of human ER-exit sites: requirements for the localisation of Sec16 to transitional ER. J Cell Sci 122, 2924–2934. doi: 10.1242/jcs.044032.

Asano, J, Tada, H, Onai, N, Sato, T, Horie, Y, Fujimoto, Y, et al. (2010). Nucleotide oligomerization binding domain-like receptor signaling enhances dendritic cell-mediated cross-priming in vivo. J Immunol 184, 736–745. doi: 10.4049/jimmunol.0900726.

Alloatti, A, Kotsias, F, Pauwels, AM, Carpier, JM, Jouve, M, Timmerman, E, et al. (2015). Toll-like Receptor 4 Engagement on Dendritic Cells Restrains Phago-Lysosome Fusion and Promotes Cross-Presentation of Antigens. Immunity 43, 1087–1100. doi: 10.1016/j.immuni.2015.11.006.

Niederberger, E, Moser, CV, Kynast, KL, and Geisslinger, G (2013). The non-canonical IkappaB kinases IKKepsilon and TBK1 as potential targets for the development of novel therapeutic drugs. Curr Mol Med 13, 1089–1097.

Reilly, SM, Chiang, SH, Decker, SJ, Chang, L, Uhm, M, Larsen, MJ, et al. (2013). An inhibitor of the protein kinases TBK1 and IKK-varepsilon improves obesity-related metabolic dysfunctions in mice. Nat Med 19, 313–321. doi: 10.1038/nm.3082.

Barrett, JC, Hansoul, S, Nicolae, DL, Cho, JH, Duerr, RH, Rioux, JD, et al. (2008). Genome-wide association defines more than 30 distinct susceptibility loci for Crohn’s disease. Nat Genet 40, 955–962. doi: 10.1038/ng.175.

Chen, JL, Stewart-Jones, G, Bossi, G, Lissin, NM, Wooldridge, L, Choi, EM, et al. (2005). Structural and kinetic basis for heightened immunogenicity of T cell vaccines. J Exp Med 201, 1243–1255. doi: 10.1084/jem.20042323.

Chu-Ping, M, Slaughter, CA, and DeMartino, GN (1992). Purification and characterization of a protein inhibitor of the 20S proteasome (macropain). Biochim Biophys Acta 1119, 303–311.

McCutchen-Maloney, SL, Matsuda, K, Shimbara, N, Binns, DD, Tanaka, K, Slaughter, CA, et al. (2000). cDNA cloning, expression, and functional characterization of PI31, a proline-rich inhibitor of the proteasome. J Biol Chem 275, 18557–18565. doi: 10.1074/jbc.M001697200.

Kirk, R, Laman, H, Knowles, PP, Murray-Rust, J, Lomonosov, M, Meziane el, K, et al. (2008). Structure of a conserved dimerization domain within the F-box protein Fbxo7 and the PI31 proteasome inhibitor. J Biol Chem 283, 22325–22335. doi: 10.1074/jbc.M709900200.

Bader, M, Benjamin, S, Wapinski, OL, Smith, DM, Goldberg, AL, and Steller, H (2011). A conserved F box regulatory complex controls proteasome activity in Drosophila. Cell 145, 371–382. doi: 10.1016/j.cell.2011.03.021.

Li, X, Thompson, D, Kumar, B, and DeMartino, GN (2014). Molecular and cellular roles of PI31 (PSMF1) protein in regulation of proteasome function. J Biol Chem 289, 17392–17405. doi: 10.1074/jbc.M114.561183.

Franchi, L, McDonald, C, Kanneganti, TD, Amer, A, and Nunez, G (2006). Nucleotide-binding oligomerization domain-like receptors: intracellular pattern recognition molecules for pathogen detection and host defense. J Immunol 177, 3507–3513.

Abbott, DW, Wilkins, A, Asara, JM, and Cantley, LC (2004). The Crohn’s disease protein, NOD2, requires RIP2 in order to induce ubiquitinylation of a novel site on NEMO. Curr Biol 14, 2217–2227. doi: 10.1016/j.cub.2004.12.032.

Yang, S, Tamai, R, Akashi, S, Takeuchi, O, Akira, S, Sugawara, S, et al. (2001). Synergistic effect of muramyldipeptide with lipopolysaccharide or lipoteichoic acid to induce inflammatory cytokines in human monocytic cells in culture. Infect Immun 69, 2045–2053. doi: 10.1128/IAI.69.4.2045-2053.2001.

Uehara, A, Yang, S, Fujimoto, Y, Fukase, K, Kusumoto, S, Shibata, K, et al. (2005). Muramyldipeptide and diaminopimelic acid-containing desmuramylpeptides in combination with chemically synthesized Toll-like receptor agonists synergistically induced production of interleukin-8 in a NOD2- and NOD1-dependent manner, respectively, in human monocytic cells in culture. Cell Microbiol 7, 53–61. doi: 10.1111/j.1462-5822.2004.00433.x.

Abbott, DW, Yang, Y, Hutti, JE, Madhavarapu, S, Kelliher, MA, and Cantley, LC (2007). Coordinated regulation of Toll-like receptor and NOD2 signaling by K63-linked polyubiquitin chains. Mol Cell Biol 27, 6012–6025. doi: 10.1128/MCB.00270-07.

Pomerantz, JL, and Baltimore, D (1999). NF-kappaB activation by a signaling complex containing TRAF2, TANK and TBK1, a novel IKK-related kinase. EMBO J 18, 6694–6704. doi: 10.1093/emboj/18.23.6694.

Delamarre, L, Pack, M, Chang, H, Mellman, I, and Trombetta, ES (2005). Differential lysosomal proteolysis in antigen-presenting cells determines antigen fate. Science 307, 1630–1634. doi: 10.1126/science.1108003.

Savina, A, Jancic, C, Hugues, S, Guermonprez, P, Vargas, P, Moura, IC, et al. (2006). NOX2 controls phagosomal pH to regulate antigen processing during crosspresentation by dendritic cells. Cell 126, 205–218. doi: 10.1016/j.cell.2006.05.035.

Gil-Torregrosa, BC, Lennon-Dumenil, AM, Kessler, B, Guermonprez, P, Ploegh, HL, Fruci, D, et al. (2004). Control of cross-presentation during dendritic cell maturation. Eur J Immunol 34, 398–407. doi: 10.1002/eji.200324508.

Wagner, CS, and Cresswell, P (2012). TLR and nucleotide-binding oligomerization domain-like receptor signals differentially regulate exogenous antigen presentation. J Immunol 188, 686–693. doi: 10.4049/jimmunol.1102214.

Samie, M, and Cresswell, P (2015). The transcription factor TFEB acts as a molecular switch that regulates exogenous antigen-presentation pathways. Nat Immunol 16, 729–736. doi: 10.1038/ni.3196.

Watson, P, Townley, AK, Koka, P, Palmer, KJ, and Stephens, DJ (2006). Sec16 defines endoplasmic reticulum exit sites and is required for secretory cargo export in mammalian cells. Traffic 7, 1678–1687. doi: 10.1111/j.1600-0854.2006.00493.x.

Luo, Y, de Lange, KM, Jostins, L, Moutsianas, L, Randall, J, Kennedy, NA, et al. (2017). Exploring the genetic architecture of inflammatory bowel disease by whole-genome sequencing identifies association at ADCY7. Nat Genet 49, 186–192. doi: 10.1038/ng.3761.

Corridoni, D, Chapman, T, Ambrose, T, and Simmons, A (2018). Emerging Mechanisms of Innate Immunity and Their Translational Potential in Inflammatory Bowel Disease. Front Med (Lausanne) 5, 32. doi: 10.3389/fmed.2018.00032.

Seiderer, J, Brand, S, Herrmann, KA, Schnitzler, F, Hatz, R, Crispin, A, et al. (2006). Predictive value of the CARD15 variant 1007fs for the diagnosis of intestinal stenoses and the need for surgery in Crohn’s disease in clinical practice: results of a prospective study. Inflamm Bowel Dis 12, 1114–1121. doi: 10.1097/01.mib.0000235836.32176.5e.

Jolles, P, Migliore-Samour, D, Maral, R, Floc’h, F, and Werner, GH (1975). Low molecular weight water-soluble peptidoglycans as adjuvants and immunostimulants. Z Immunitatsforsch Exp Klin Immunol 149, 331–340.

