## Supplementary Material for "NOD2 and TLR2 Signal via TBK1 and PI31 to Direct Cross-presentation and CD8 T Cell Responses"

**Supplementary Materials**

**A**

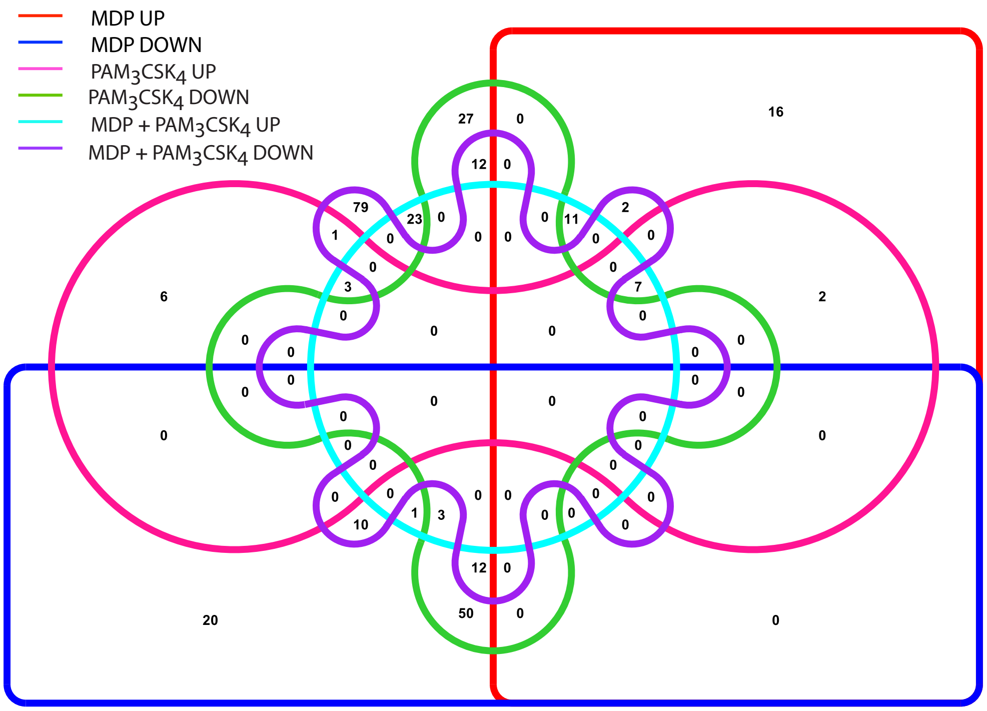


**Figure S1. Number of proteins detected as up and down regulated by phosphoproteomic analysis. (A)** Quantitative phospho-proteomic in DCs stimulated for 30 min with MDP (10 µg/ml), $PAM_3CSK_4$ (1 µg/ml) and combined MDP +$PAM_3CSK_4$, followed by phosphoenrichment (PE) and generation of phosphopeptides for LC-MS/MS. Venn diagram showing number of up- and down-regulated proteins detected following MDP, $Pam_3CSK_4$ and dual stimulation. 134 proteins showed differential regulation when stimulated with MDP (38 up-regulated, 96 down-regulated), 123 when stimulated with Pam3CSK4 (19 up-regulated, 104 down- regulated) and 164 when stimulated with both MDP and Pam3CSK4 (48 up-regulated, 116 down-regulated).

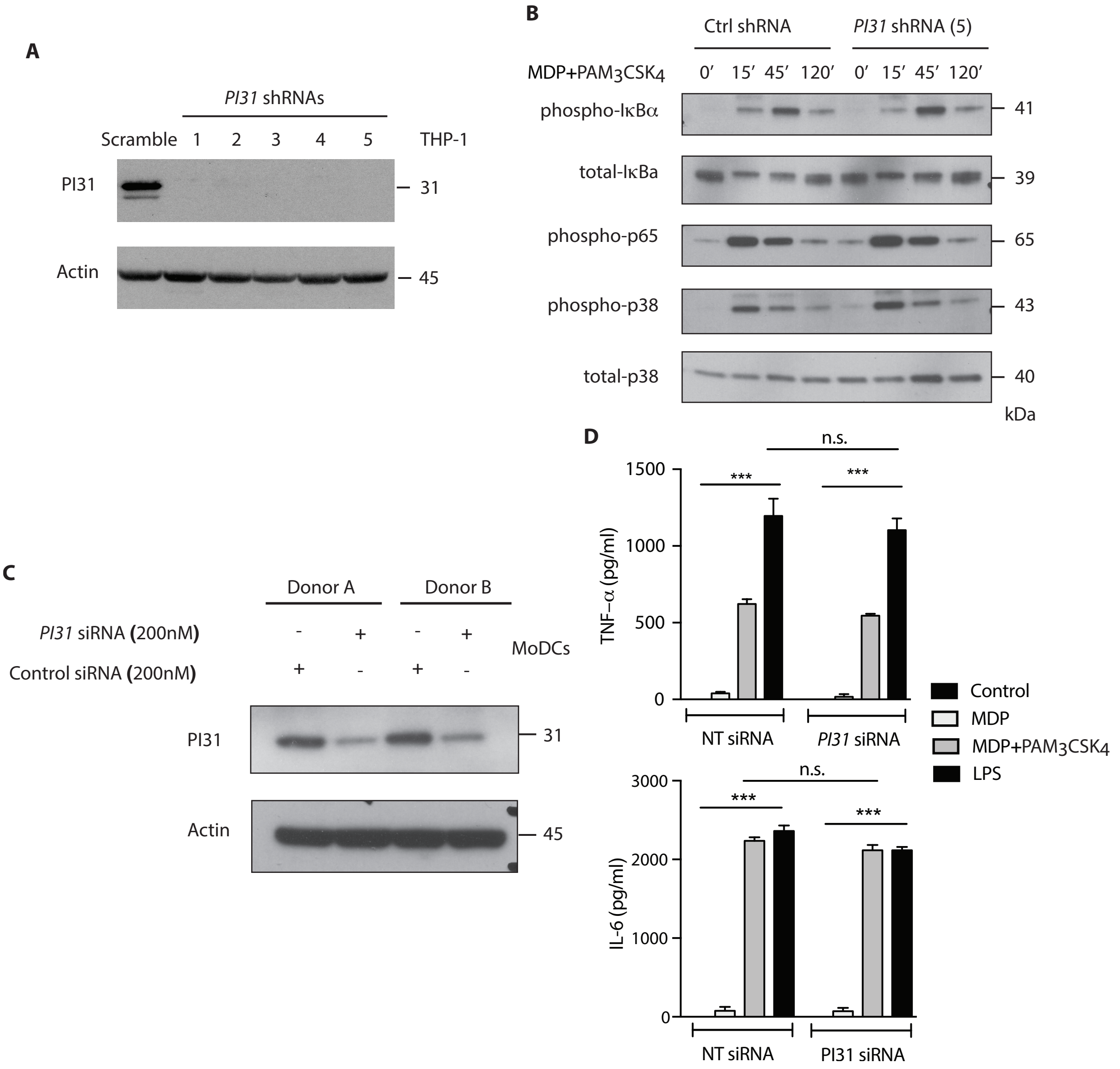


**Figure S2. PI31 does not affect NF-κB and MAPK activation or cytokine secretion on NOD2/TLR2 triggering.** (**A**) THP1 cells were transduced with control or *PI31*-targeting lentiviral shRNAs and analysed for PI31 expression by western blot. (**B**) Immunoblot analysis of phospho-Ikba, total- Ikba, phospho-p65, phospho-p38 and total-p38 in THP1 cells transduced with Ctrl or PI31 shRNAs and stimulated with MDP (10 μg/ml) and $PAM_3CSK_4$ (1 μg/ml) for 15’ 45’ and 120 min. Immunoblots data represent one experiment representative of two separate experiments. (**C**) Primary DCs cells were transfected with Ctrl non-targeting or *PI31* targeting siRNAs. PI31 levels were analyzed by western blot. (**D**) ELISA of TNF-α and IL-6 released from DCs transfected as in (**C**) and stimulated for 24 hours with MDP (10 μg/ml), $PAM_3CSK_4$ (1 μg/ml), their combination or LPS (10ng/ml). Data represent mean ± s.e.m ($n$ = 8), ***$P$<0.0001 Student’s $t$ test. Data are from two separate experiments.

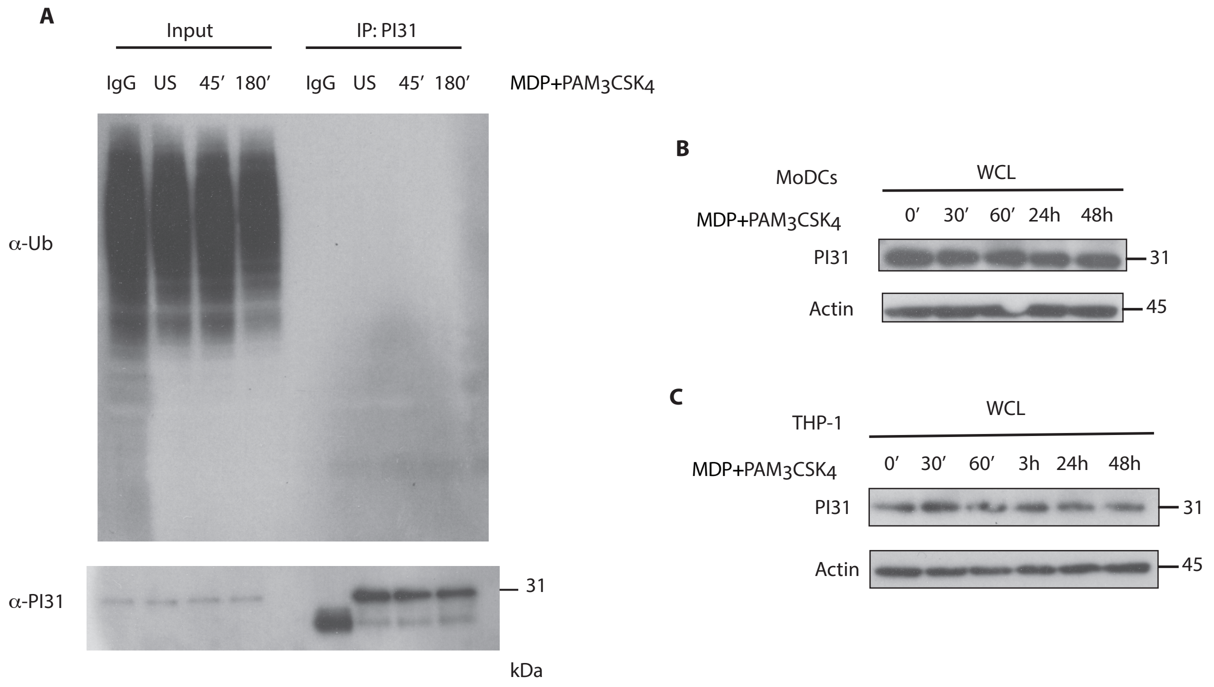


**Figure S3. PI31 is not polyubiquitinated and degraded following NOD2/TLR2 triggering.** (**A**) PI31 was immunoprecipitated from DC lysates stimulated with MDP (10 µg/ml) and $PAM_3CSK_4$ (1 µg/ml) for 45 and 180 min. Immunoblot using anti-FK2 antibody is shown (upper panel) and with anti-PI31 (lower panel). Immunoblots data from one experiment representative of two separate experiments. Primary DCs (**B**) and THP1 (**C**) cells were stimulated with MDP (10 µg/ml) and $PAM_3CSK_4$ (1 µg/ml) for 30, 60 minutes for either 24 or 48 hrs. PI31 levels were analysed by immunoblot analysis. Immunoblots data from one experiment representative of three separate experiments.

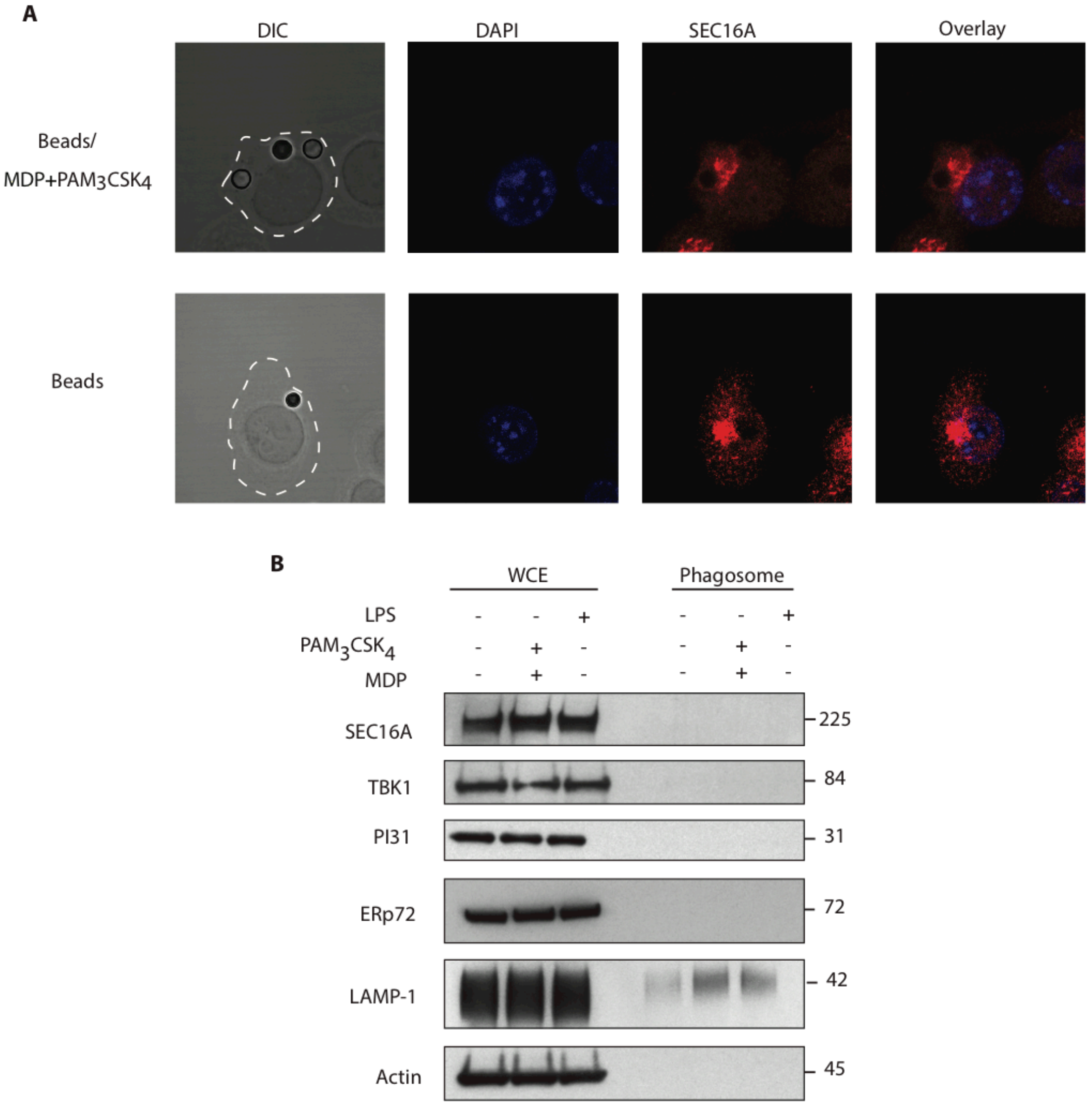


**Figure S4. SEC16A, PI31 and TBK1 are not enriched in phagosomes containing NOD2 and TLR2 ligands.** (**A**) BMDCs were exposed to unconjugated beads or beads conjugated to MDP and $PAM_3CSK_4$. Representative SEC16A staining analyzed by confocal microscopy. (**B**) DC whole cell lysates (WCE) and phagosomal lysates were immunoblotted for Sec16A, TBK1, PI31, Erp72 and LAMP1. Immunoblot for actin is shown in the lower panel. Data from one experiment representative of three separate experiments.
